# Single-cell multiomics reveals archetypal regulatory programs shared across CD4 and CD8 T cell subsets in viral infection

**DOI:** 10.1101/2025.09.08.675014

**Authors:** Sarah K. Walker, Joris van der Veeken, Alexander Rudensky, Yuri Pritykin

## Abstract

T cells protect against pathogens and tumors and can differentiate into various functionally distinct subsets. While each subset exhibits a characteristic epigenomic and transcriptional profile, essential immune functions such as self-renewal, expansion, cytokine production, and cytotoxicity are common to multiple subsets and therefore must be programmed via shared regulatory mechanisms. To uncover these regulatory “archetypes”, we integrated a new single-cell multiomic (scATAC+RNA-seq) dataset for CD4 and CD8 T cells responding to acute and chronic viral infection with a comprehensive compendium of bulk ATAC-seq data. Our analysis revealed that T cell identity is governed by a modular architecture, where distinct transcription factors drive the reuse of regulatory archetypes across T cell subsets and lineages. Notably, translationally important progenitor Tcf1^+^ CD8 T cells, critical for sustaining CD8 T cell responses and present in both acute and chronic infection, exhibited a composite regulatory state combining a CD8 T cell exhaustion program and, unexpectedly, a follicular helper CD4 T cell program. In sum, this resource will aid mechanistic dissection of adaptive immunity and immunotherapy design, and the framework will be broadly applicable across biological systems.

**Highlights:**

– Single-cell multiomics and ATAC-seq atlas reveal T cell regulatory archetypes
– A unified progenitor Tcf1^+^ CD8 T cell population is shared by acute and chronic infection
– Progenitor CD8 T cell identity combines CD8 exhaustion and CD4 Tfh regulatory programs
– A Tfh-like differentiation gradient underlies heterogeneity in progenitor CD8 T cells

## Introduction

T cells coordinate systemic immunity and protect against infections and tumors. Over their lifespan and in response to antigen stimulation, T cells differentiate into distinct lineages and functional states, each characterized by unique epigenomic and transcriptional profiles.^1–4^ However, many core immune functions, such as self-renewal, expansion, cytotoxicity, production of and response to particular cytokines, and tissue migration, are not unique to a specific T cell subset, but instead recur across subsets. Therefore, these recurring functions must be driven by common underlying regulatory programs. Indeed, flow cytometry analysis has revealed that activity of many key functional molecules, surface markers and transcription factors (TFs) spans multiple T cell populations, including across CD4 and CD8 lineages. For example, interferon-*γ* (IFNg) production is a hallmark of both effector CD8 T cell and CD4 Th1 cell responses; adhesion molecule L-selectin (CD62L) marks both naive and central memory T cells; both follicular helper CD4 T cells (Tfh) and exhausted CD8 T cells express the inhibitory receptor programmed death 1 (PD-1) and the transcription factor TOX; and the chemokine receptor CXCR5 is shared between Tfh and progenitor exhausted CD8 T cells.^1, 3–10^ A central challenge in T cell biology is to dissect the shared and context-specific “archetypal” regulatory programs that govern both overlapping and specialized T cell functions. Achieving this requires comprehensive and high-resolution approaches capable of linking gene regulatory potential with transcriptional output. While flow cytometry with a limited set of markers is effective for identifying well-known populations, it restricts the unbiased comparison of similarities and differences of discrete and continuous T cell phenotypes across models. Single-cell multiomic technologies that simultaneously profile chromatin accessibility and gene expression at the genome-wide level now provide an opportunity to resolve the archetypal regulatory programs systematically across heterogeneous T cell populations.^11–18^

The lymphocytic choriomeningitis virus (LCMV) infection model offers a powerful experimental system to investigate such regulatory programs across divergent immune responses. Profiling T cells at day 7 post-infection with either the Armstrong strain (acute) or Clone 13 (chronic) captures a critical point of divergence between viral clearance and persistent infection leading to exhaustion.^7, 8, 19, 20^ Although both infections elicit overlapping T cell subsets, they differ markedly in antigen persistence and immune outcomes, providing a tractable framework to study shared versus context-specific features of T cell regulation. Including both CD4 and CD8 T cells in this analysis helps fill a significant gap in prior genomic studies of the LCMV model, which have provided foundational insights into CD8 T cells while leaving CD4 subsets and especially their relationship with CD8 responses relatively understudied.^6, 21–24^

A particularly important and intensively studied population is the TCF1^+^ progenitor CD8 T cell subset, which is essential for sustaining responses in chronic infection and cancer, and for enabling effective T cell reactivation in cancer immunotherapies.^7–9, 25–32^ While this population, often termed “progenitor exhausted” cells, has been extensively studied in chronic contexts, it has recently been observed in acute responses and shares features with memory precursor cells.^33, 34^ These cells have also been termed “stem-like” or “memory-like,” reflecting their capacity for longterm persistence and functional flexibility. The precise relationship between progenitor-like populations in acute versus chronic contexts has been an active area of investigation. Some studies have described distinct precursor populations in each setting,^19, 35–37^ whereas others have observed a shared progenitor state across contexts.^32, 38–40^ A systematic, high-resolution analysis is needed to reconcile these observations and provide a unified epigenomic definition of this critical cell population. Furthermore, prior studies have noted that progenitor CD8 T cells share molecular markers and functional properties with CD4 Tfh cells.^10, 41^ Tfh cells are a specialized subset of CD4 T cells that provide essential help to B cells during the germinal center response, promoting antibody production and affinity maturation.^10^ They are characterized by expression of CXCR5, PD1, TOX, TCF1, and a “master regulator” factor BCL6. Progenitor exhausted CD8 T cells, initially defined as a CXCR5^+^ subset within the exhausted CD8 T cell pool, also express PD1 and TOX and have enriched expression of TCF1 and BCL6. Functionally, they show similarities with Tfh cells: both are defined by sustained TCR stimulation and limited proliferation without terminal effector differentiation.^8, 10^ However, a key distinction is that progenitor CD8 T cells exhibit persistence and retain both self-renewal capacity and stem-like behavior, features not observed in Tfh cells. Despite these parallels, the extent of the regulatory and functional similarity between progenitor CD8 T cells and CD4 Tfh cells remains incompletely understood. A systematic comparison of their epigenomic and transcriptomic features in the context of infection response provides an opportunity to define common regulatory modules and clarify their distinct features.

In this study, we performed a joint single-cell chromatin accessibility and transcriptomic profiling (scATAC + RNA-seq) of T cells responding to viral infection. Chromatin accessibility reflects the regulatory potential of the cell and provides direct insight into TF activity.^16, 32, 35, 36, 42–44^ However, in contrast to gene expression analysis using scRNA-seq, which benefits from reliable public reference gene annotations, chromatin accessibility is more plastic and cell type-specific, so scATAC-seq analysis requires building a cell type-specific reference set of accessible regions, which is not readily available for T cells. To address this, we compiled a compendium of 285 bulk ATAC-seq profiles collected from 17 different publications and built a comprehensive reference atlas of chromatin accessibility regions in CD4 and CD8 T cells. We developed a new computational pipeline that leverages this atlas and enables high-resolution, biologically interpretable analysis of T cell scATAC-seq and multiomic scATAC+RNA-seq data. We identified all major CD4 and CD8 T cell functional states in our single-cell multiomic data, including naive, memory, effector, regulatory, Tfh, exhausted and progenitor populations, each present in both LCMV Armstrong and Clone 13 responses, albeit at different frequencies. In our data, and by improved reanalysis of published datasets, we demonstrated that Tcf1^+^ progenitor CD8 T cells from acute and chronic responses represent a shared, transcriptomically and epigenomically similar population. Furthermore, we uncovered striking similarity in chromatin accessibility and gene expression between progenitor CD8 T cells and follicular helper CD4 T cells, suggesting the adoption of shared regulatory programs by these two populations. To systematically decompose the regulatory landscape of T cells, we applied archetypal analysis to chromatin accessibility profiles. This approach identified a small number of “extreme” states, or *archetypes*, that capture the major axes of variation in the data. These archetypes recapitulated key biological programs, both unique to specific T cell populations, and shared between populations, such as cytotoxicity, exhaustion, quiescence, and Tfh-like regulation. TF motif analysis implicated potential drivers of these regulatory programs; it was further validated using allele-specific analysis, leveraging natural genetic polymorphisms in our data, which was generated in F1 hybrid of two mouse strains. Remarkably, unlike multiple other functional T cell states, Tcf1^+^ progenitor CD8 T cells did not align with a single archetype, but instead exhibited a composite regulatory identity combining exhaustion- and Tfh-associated archetypes. This decomposition provides a mechanistic explanation for the shared features of progenitor CD8 T cells and CD4 Tfh cells and reveals that progenitor phenotype may emerge from the modular reuse of regulatory programs active in other T cell contexts.

Together, our work introduces a powerful framework for high-resolution, interpretable singlecell multi-omic analysis and reveals shared regulatory logic across T cell populations responding to viral infection, linking progenitor CD8 and helper CD4 T cell programs. By integrating computational analysis with comprehensive experimental profiling of both CD4 and CD8 T cells, we provide a valuable resource and conceptual advance for understanding the transcriptional control of adaptive immune responses.

## Results

### ATAC-seq compendium enables robust analysis of scATAC+RNA-seq data for T cells in LCMV infection

We used single-cell genomics to systematically characterize both discrete and continuous phenotypes and regulatory states of CD4 and CD8 T cells during the early stages of the viral response. We generated single-cell multiomic datasets capturing both chromatin accessibility (scATAC-seq) and gene expression (scRNA-seq) in splenic T cells at day 7 post-infection with either LCMV Armstrong (Arm) or LCMV Clone 13 (Cl13). To enhance the resolution and reproducibility of chromatin accessibility measurements, we also generated standalone scATAC-seq data for the same cell populations (**Fig. 1A, S1, Methods**), yielding a total of 31,662 cells. These experiments were performed in F1 hybrid mice generated by crossing the laboratory C57BL/6 (B6) and wild-derived CAST/EiJ (Cast) strains, allowing both allele-agnostic analysis, which forms the primary focus of this study, and allele-specific analysis of gene expression regulation (described below).

**Fig. 1.**
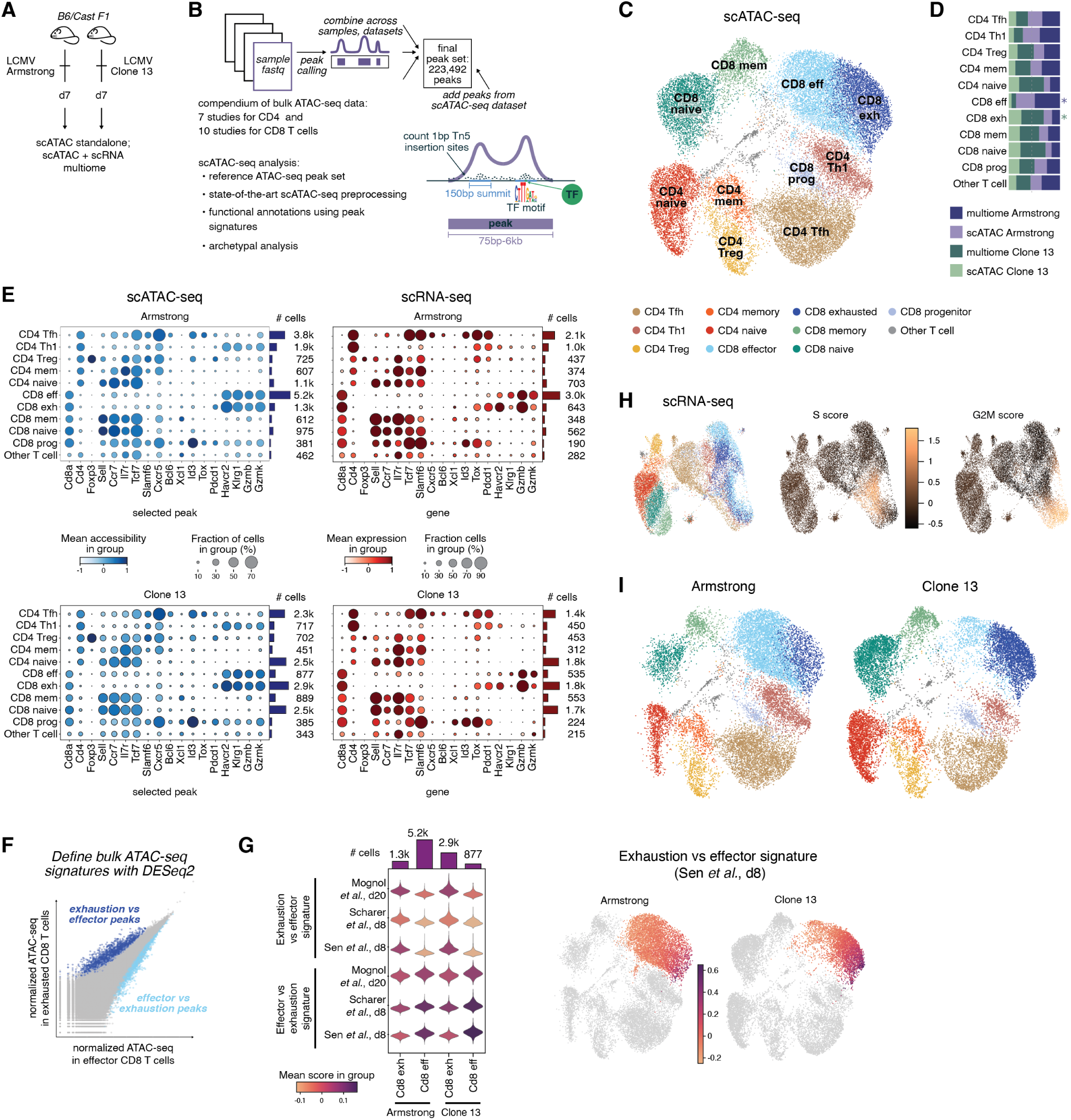
Early T cell responses to acute and chronic infection share the same functional cell states. **(A)** Experimental setup: splenic T cells from F1 hybrid C57BL/6J *×* CAST/EiJ mice infected by either Armstrong clone (Arm, acute infection) or Clone 13 (Cl13, chronic infection) of Lymphocytic choriomeningitis virus (LCMV) were profiled either with scATAC-seq or multi-omic scATAC+scRNA-seq 7 days (d7) post infection. **(B)** Summary and schematic of the T cell scATAC-seq analysis. For the T cell ATAC-seq peak atlas, peaks were called using publicly available bulk ATAC-seq datasets for T cells from both CD4 and CD8 lineages, and combined with peaks from the new scATAC-seq dataset. Key aspects of our scATAC-seq data analysis pipeline highlighted. **(C)** Uniform manifold approximation and projection (UMAP) of scATAC-seq data with T cell functional annotations. **(D)** Barplot showing the distribution of functional cell states across the four samples (LCMV Arm and Cl13 for both standalone scATAC-seq and scATAC-seq from the multiome profiling). Asterisks, significant enrichments (hypergeometric test, odds ratio *>* 2, *p <* 1.3*e −* 70). **(E)** Dotplots showing normalized chromatin accessibility levels at selected peaks for marker genes (left) and normalized expression of selected marker genes (right) in different T cell functional states, for Arm (top) and Cl13 (bottom) cells. Dot size, fraction of cells with normalized accessibility or expression greater than 0; dot color, mean normalized accessibility or expression in that cell state. Barplots on the right side show the number of cells per cell state. **(F)** Example of how epigenomic signatures are generated using DESeq2 for bulk ATAC-seq data for two conditions from the same dataset. Peak signatures are formed from differentially accessible peaks in either direction (LFC *>* 1 or LFC *< −*1, FDR *≤* 0.05). The scATAC-seq data is then scored for enrichment of these peak signatures. **(G)** *Left*, violin plots showing distributions of the epigenomic signature scores in effector and exhausted CD8 T cells, separately in Arm and Cl13 samples. The signatures are for CD8 T cells at d8 in LCMV infection from multiple studies.^45–47^ In each dataset, peaks significantly more accessible in exhausted than in effector cells (exhaustion vs. effector signature) and in effector than in exhausted cells (effector vs. exhaustion signature) were defined, and then scored based on accessibility of these peaks in our single-cell data. Top margin, the barplot of the number of cells in each group. All scores were significantly different between effectors and exhausted groups (FDR *<* 1*e −* 46, two-sided Mann-Whitney U test). *Right*, UMAP plots showing the scores of the exhaustion vs. effector signature derived from Sen *et al.*^47^ in cells annotated as either effector or exhausted CD8 T cells, separately for the Arm and Cl13 responses. **(H)** *Left*, UMAP of the scRNA-seq modality of the dataset showing T cell functional states annotated using the scATAC-seq analysis. *Middle and right*, same UMAP showing cell cycle gene signature scores (S phase, middle; G2M phase, right). **(I)** UMAP with the functional T cell state annotations separately for cells from Arm and Cl13 infection.

We developed a new computational pipeline to analyze this multiomic dataset, integrating best practices from previous methods and building a new chromatin accessibility reference for T cells (**Methods**). We systematically defined a comprehensive set of accessible regions in T cells using publicly available bulk ATAC-seq datasets. Building on our prior resource for CD8 T cells,^32^ we assembled and uniformly reprocessed a compendium of 285 ATAC-seq profiles covering various functional states of CD8 and CD4 T cells across 17 studies (**Fig. 1B, Table S1, Methods**). This enabled the construction of a high-resolution atlas of chromatin accessibility peaks for T cells across multiple mouse models. To further enhance the coverage of the T cell regulatory landscape in T cells, we extended this atlas with peaks newly identified from our scATAC-seq data. The resulting union set of 223,492 peaks served as the basis for all subsequent analyses (**Table S2**).

Previous studies have shown that chromatin accessibility profiles can better distinguish cellular phenotypes than transcriptomic profiles, and thus can better reflect cell identity and function.^18, 48, 49^ Therefore, we used our scATAC-seq data as the primary modality for comprehensive functional annotation. Following normalization, dimensionality reduction, and integration of the standalone scATAC-seq and the scATAC-seq component of the multiome, we performed clustering to define and annotate reproducible subpopulations of CD4 and CD8 T cells (**Fig. 1C–E, S2**). We annotated these clusters using established marker genes, based on both their gene expression and the activity of associated chromatin accessibility peaks (**Fig. 1E, S2D, S3, Table S3, Table S4, Table S5, Methods**). We identified all major CD4 and CD8 T cell functional states, including naive, memory, effector, regulatory, Tfh, exhausted and progenitor populations. In particular, we observed a continuum of actively responding CD8 T cells which we classified into hyperactivated cells in the early state of exhaustion (referred to as exhausted), more frequent in Cl13 response and associated with higher activity of inhibitory and exhaustion marker genes *Pdcd1*, *Tox*, *Havcr2* (*Tim3* ), *Lag3* and *Tigit* and elevated cytotoxicity as indicated by overexpression of *Gzmb*, *Nkg7* and *Prf1*, and the remaining effector cells more frequent in the Arm response (**Fig. 1C–E, S3**). To further support our annotations, we derived functional epigenomic signatures from the bulk ATAC-seq compendium (**Fig. 1F, Table S3, Methods**). We identified peaks distinguishing known T cell subsets and scored these peak signatures in scATAC-seq data, assigning higher scores to cells with greater accessibility at subset-specific peaks. This analysis further supported our annotations by capturing key contrasts, e.g. between naive and memory T cells (**Fig. S2F,G**) and between effector and exhausted CD8 T cells (**Fig. 1G, S2H**). Furthermore, all T cell subpopulations were observed in both LCMV Arm and Cl13 in both standalone scATAC-seq and the multiome scATAC-seq, demonstrating reproducibility across technologies (**Fig. 1D,E**). Thus, all major functional subsets of CD4 and CD8 T cells are reproducibly present in both LCMV Arm and Cl13 responses, albeit in different frequencies.

We observed that applying standard scATAC-seq analysis pipelines can sometimes group cell populations in ways that are challenging to interpret biologically (**Fig. S4, S5, Methods**). Furthermore, clustering based on the scRNA-seq modality grouped transcriptionally similar but functionally distinct populations, such as naive and memory T cells, or proliferating CD4 and CD8 T cells (**Fig. 1H, S6**). These observations are consistent with prior work suggesting chromatin accessibility can be a more stable marker of cell identity than transcriptomics, and highlight the power of the improved resolution of our scATAC-seq pipeline integrating single-cell data with a comprehensive reference atlas.

In summary, our computational pipeline leveraging a new comprehensive T cell ATAC-seq compendium to analyze our new high-resolution multiomic dataset enabled accurate annotation of T cell functional and differentiation states in early LCMV infection. We found that all major CD4 and CD8 T cell states are present in both acute and chronic responses but in different proportions. The presence of exhausted-like CD8 T cells in acute infection challenges the traditional view that exhaustion is restricted to chronic infection or cancer. However, this observation is consistent with recent studies reporting transcriptional and epigenetic features of exhaustion emerging early and transiently in acute responses before viral clearance.^38, 50^ Our findings reinforce this emerging perspective by demonstrating that exhaustion-associated states are not exclusive to persistent antigen exposure but may be part of a broader T cell activation and differentiation landscape.

### Shared progenitor Tcf1^+^ CD8 T cell state in early LCMV Armstrong and clone 13 responses

We have previously reported a subpopulation of Tcf1^+^ progenitor-like CD8 T cells shared between acute and chronic responses, using scATAC-seq and scRNA-seq data analysis in LCMV Arm and Cl13, and further supported this finding through reanalysis of multiple previously published scRNA-seq datasets.^32^ Recent functional studies, including adoptive transfer experiments, have further confirmed that the progenitor population is shared between acute and chronic infection.^38–40^ Nevertheless, earlier studies described stem-like progenitor exhausted Cxcr5^+^ Tcf1^+^ CD8 T cells exclusively in chronic infection and subsequently cancer, while Il7r^+^ Klrg1^−^ memory precursor cells were described in acute infection.^33, 34^ Some reports, including using high-throughput genomic analysis, proposed that these represent distinct subpopulations.^19, 35–37^ To reconcile these results, we further explored these CD8 T cell phenotypes using multiomic single-cell data and our refined computational pipeline.

Our new analysis supports the presence of epigenomically and transcriptionally similar population of memory-like progenitor or precursor Tcf1^+^ CD8 T cells in both LCMV Arm and Cl13 (**Fig. 2A,B**). This population was reproducible between standalone and multiome scATAC-seq (**Fig. 1D**) and showed enrichment of transcriptional activity of genes associated with both progenitor exhausted and memory precursor cells, such as *Cxcr5*, *Slam6*, *Tox*, *Tcf7*, *Id3* and *Il7r*, even when analyzed separately by infection type (**Fig. 1E, 2B**). Moreover, cells in this population were consistently enriched for the progenitor ATAC-seq peak signatures derived from two different studies, again reproducibly across both LCMV Arm and Cl13 (**Fig. 2A**). Taken together, these observations reinforce the existence of an epigenomically and transcriptionally similar population of stem-like progenitor Tcf1^+^ CD8 T cell state in both acute and chronic viral infection.

**Fig. 2.**
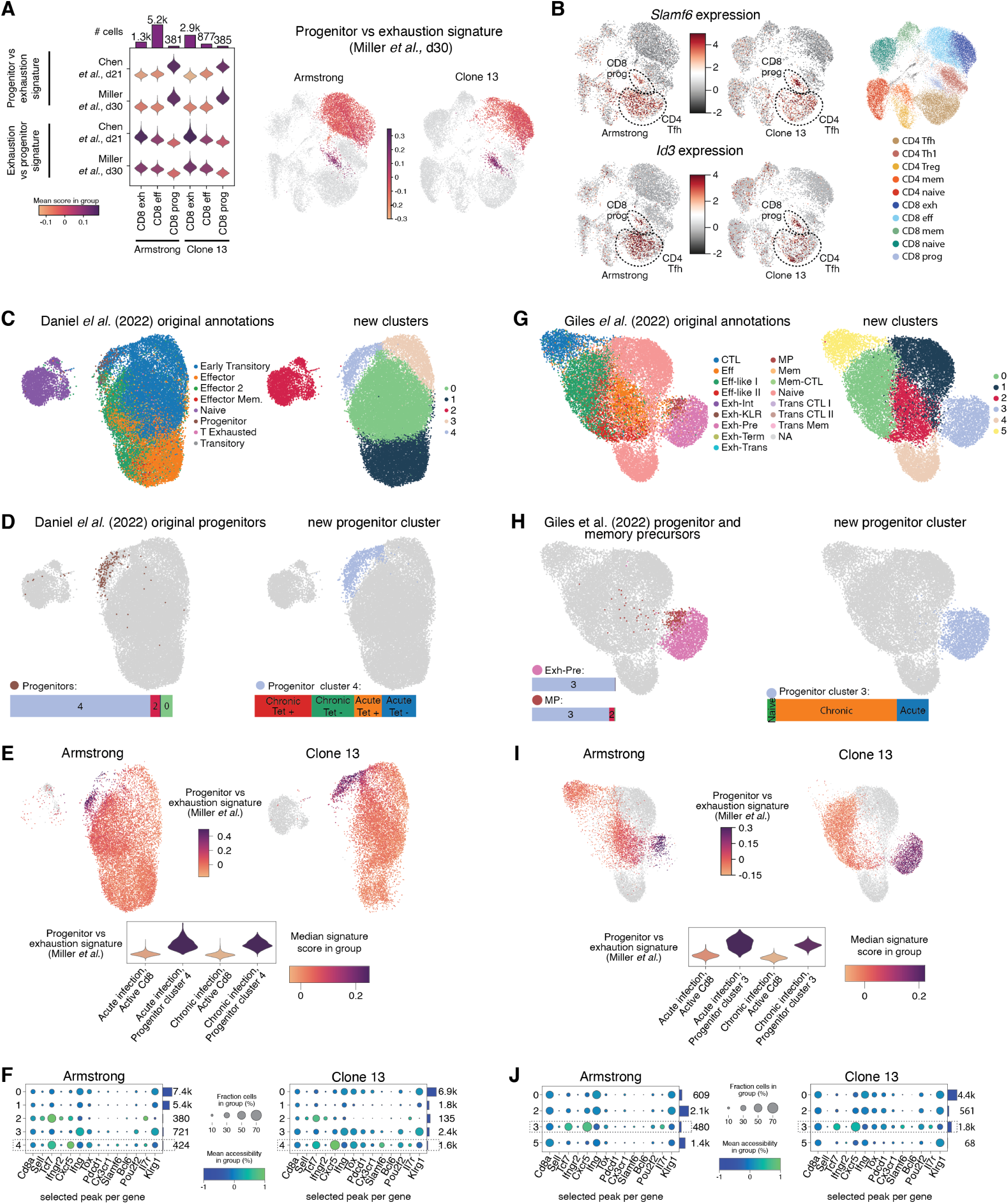
Shared progenitor Tcf1+ CD8 T cell state in early response to acute and chronic viral infection. **(A)** *Left*, violin plots of the epigenomic signature scores comparing progenitor vs. exhaustion T cell states (similar to Fig. 1G). The signatures were defined using data from progenitor and terminally exhausted CD8 T cells at d21 in response to melanoma^51^ or at d30 in response to LCMV Cl13 infection.^52^ All scores were significantly different between progenitor and effector or dysfunctional CD8 T cell groups (FDR *<* 5*e −* 52, two-sided Mann-Whitney U test). *Right*, UMAP plots showing the epigenomic signature scores of progenitor vs. terminal exhaustion from Miller *et al.*^52^ in cells annotated as either progenitor, effector, or exhausted CD8 T cells in our data. **(B)** UMAP plots showing normalized scRNA-seq gene expression for the two progenitor marker genes, *Id3* and *Slamf6*, separately for Armstrong and Clone 13 cells. **(C-F)** Reanalysis of the scATAC-seq data for CD8 T cells at day 8 in response to LCMV infection from Daniel *et al.*^35^ The dataset was preprocessed and quantified using our T cell ATAC-seq peak atlas and then analyzed using our computational pipeline. Cluster 4 in the reanalysis are progenitor cells. **(C)** UMAP plots with the cell state annotations from the publication^35^ (left) and our newly defined clusters (right). **(D)** UMAP showing the progenitor cells. *Left*, cells annotated as progenitors in the publication.^35^ The barplot shows the distribution of these annotations within our clusters in our reanalysis for these cells. *Right*, cells annotated as progenitors (cluster 4) in our reanalysis. The barplot shows the distribution of samples of origin for these cells. **(E)** *Top,* UMAP plots showing the score for the progenitor vs. exhaustion epigenomic signature (as in panel A), separately for the Arm (left) and Cl13 (right) samples, for all cells except naive that are shown in gray. *Bottom,* violin plot showing the progenitor vs. exhaustion score in non-naïve cells and all cluster 4 progenitor cells, separately for Arm and Cl13 infection. **(F)** Dotplots showing chromatin accessibility for selected peaks near marker genes in clusters from our reanalysis of the data, separately for the Arm (left) and Cl13 (right) samples, with the progenitor cluster 4 highlighted. The dot size, fraction of cells with normalized accessibility greater than 0; dot color, mean normalized accessibility in that cluster. Barplots on the right side show the number of cells per cluster. **(G-J)** Same as panels (B-E) but for reanalysis of a dataset for CD8 T cells at d8 post LCMV infection from Giles *et al*.^36^ Cluster 3 in this reanalysis are progenitor cells. **(G)** UMAP plots with the cell state annotations from the publication^36^ (left) and our newly defined clusters (right). **(H)** UMAP showing the progenitor cells. *Left*, cells annotated as exhausted progenitor or memory precursor cells in the publication.^36^ The barplot shows the distribution of cluster annotations in our reanalysis for these cells. *Right*, cells annotated as progenitors (cluster 3) in our reanalysis. The barplot shows the distribution of samples of origin for these cells. **(I)** *Top,* UMAP plots showing the progenitor vs. exhaustion epigenomic signature scores, separately for the Arm (left) and Cl13 (right) samples, for all cells except naive that are shown in gray. *Bottom,* violin plot showing progenitor score in active (non-naïve) cells and all cluster 3 progenitor cells, separately for acute (Arm) and chronic (Cl13) infection. **(J)** Dotplots showing chromatin accessibility for selected peaks in clusters from our reanalysis of the data, separately for the Arm (left) and Cl13 (right) samples (excluding clusters 1 and 4 that contain only naive cells, and naive cells from other clusters). The dot size, fraction of cells with normalized accessibility greater than 0; dot color, mean normalized accessibility in that cluster. Barplots on the right side show the number of cells per cluster.

**Fig. 3.**
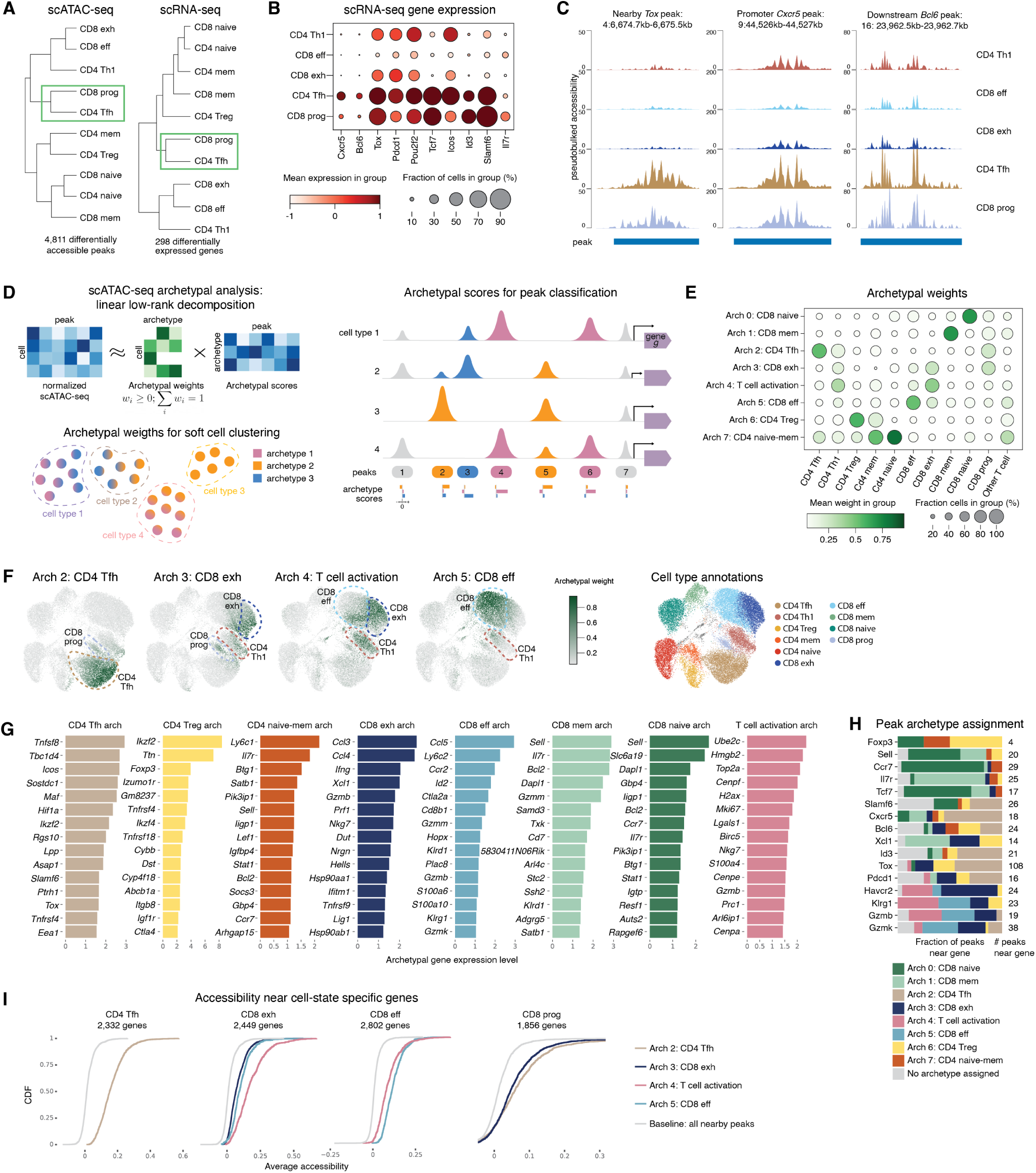
Archetypal analysis of scATAC-seq data reveals the mixed regulatory program of progenitor Tcf1^+^ CD8 T cells and their similarity with follicular helper CD4 T cells. **(A)** Hierarchical clustering of pseudobulk scATAC-seq (left) and scRNA-seq (right) profiles for T cells in different functional states. Significantly differentially accessible peaks (LFC *>* 1.5, FDR *≤* 0.05 for any cell state vs. other cells, left) and differentially expressed genes (LFC *>* 1.5, FDR *≤* 0.05 for any cell state vs. other cells, right) were used for this analysis. **(B)** Dotplot showing expression of selected Tfh and progenitor marker genes for activated T cell subsets. Dot size, fraction of cells with normalized expression greater than 0; dot color, mean normalized expression in that cell state. **(C)** Pseudobulk scATAC-seq tracks for activated T cell subsets at selected peaks (shown as horizontal blue bars at the bottom) of the genes *Tox*, *Cxcr5*, and *Bcl6*. **(D)** Schematic of the archetypal analysis of scATAC-seq data. *Top left* : archetypal analysis relies on a linear low-rank matrix decomposition of the data (Methods). *Bottom left*: archetypal analysis enables soft cell clustering via the archetypal weight matrix. Each cell can be assigned to multiple archetypes with some probability (referred to as an archetypal weight). Each archetype can be considered as a pseudo-cell averaging over cells with a similar peak accessibility distribution. *Right* : archetypal analysis assigns a score to each peak, reflecting its contribution to each chromatin accessibility archetype. For downstream analysis, each peak is uniquely assigned to the archetype with the highest score; peaks without a predominant archetype are not assigned to any (shown in gray). A gene may have multiple peaks coming from different archetypes, which may be associated with distinct programs of differential regulation across functional cell states and conditions. **(E-I)** Results of archetypal analysis of our T cell scATAC-seq data using eight archetypes (Methods). **(E)** Dotplot showing archetypal weights (probability of an archetype in a cell) for different cell states. Dot size, fraction of cells with archetypal weight greater than 0; dot color, mean archetypal weight in that cell state. **(F)** UMAP plots showing archetypal weights for four example archetypes. Cell type annotations are shown for reference. **(G)** Top protein-coding genes expressed in each scATAC-seq archetype (Methods). **(H)** Barplot of the distributions of peak assignments to archetypes for selected genes. **(I)** Analysis of cell state-specific chromatin accessibility in peaks associated with genes with cell state-specific expression, for four selected cell states: CD4 Tfh, exhausted CD8 T cells, CD8 effectors, and CD8 progenitor cells. Genes with significant differential overexpression (LFC *>* 0, FDR *≤* 0.05) in a cell state were selected. For these genes, cumulative distribution function (CDF) plots show the average accessibility of nearby peaks that are associated with a particular archetype. The gray lines indicate the distribution of the average accessibility for all peaks near a gene, used as a baseline. Only the archetypes that have significantly increased accessibility compared to the baseline are shown for each cell state (KS test, FDR *≤* 0.05 and difference in medians *≥* 0.03).

Nevertheless, two recent scATAC-seq analyses of CD8 T cells in LCMV Arm and Cl13 reported largely non-overlapping T cell subsets between the two infections, including a distinct progenitor exhausted subpopulation in LCMV Cl13 and a separate memory precursor population in LCMV Arm.^35, 36^ We hypothesized that these apparently distinct populations might represent a single underlying state whose annotation is sensitive to analytical parameters and the reference set of regulatory elements used. We reasoned that reanalyzing these important datasets using our comprehensive T cell compendium could provide a unified perspective and help reconcile the different interpretations.

To test this, we reanalyzed the Daniel *et al.* dataset^35^ using our computational pipeline and T cell ATAC-seq compendium, focusing on the day 8 timepoint to match our own day 7 analysis. We identified cluster 4 as progenitor Tcf1^+^ CD8 T cells based on known marker gene activity and enrichment of the progenitor-specific epigenomic signature, observed consistently in both LCMV Arm and Cl13 samples (**Fig. 2C–F**). This cluster included 86% of the cells originally annotated as progenitor exhausted,^35^ as well as other cells from both infections (**Fig. 2D**). A comparison suggested that our reanalysis provided a more biologically consistent annotation (**Fig. S7A,B**). Similarly, reanalysis of the scATAC-seq dataset from Giles *et al.*^36^ using our new computational pipeline identified cluster 3 as progenitors, again supported by marker gene activity and epigenomic signatures across both infections (**Fig. 2G–J**). This cluster contained 99.8% of cells originally annotated as progenitor exhausted (from Cl13) and 92% of those labeled as memory precursors (from Arm) by the authors,^36^ suggesting that the two groups represent a similar underlying progenitor state (**Fig. 2H**). Again, a more detailed comparison suggested that our updated annotations were more biologically consistent (**Fig. S7C,D**).

Together, these results demonstrate that our computational approach, leveraging a comprehensive bulk ATAC-seq reference and epigenomic signatures of functional cell states, enables more comprehensive and biologically consistent annotation of progenitor Tcf1^+^ CD8 T cells. Applied to both our new multiomic data and previously published datasets, this analysis supports the existence of a shared progenitor population in both acute and chronic viral responses.

### Progenitor Tcf1^+^ CD8 T cells and follicular helper CD4 T cells have similar chromatin accessibility and gene expression profiles

Next, we wanted to use our single-cell multiomic dataset for systematic analysis of similarities and differences of regulatory programs across CD4 and CD8 T cell lineages in response to LCMV infection. (**Fig. 3A, S8A,B**). Strikingly, antigen activation status appeared to be a stronger determinant of regulatory and phenotypic similarity than lineage identity. For example, naive and memory CD4 and CD8 T cells clustered together, while effector and exhausted CD8 T cells clustered more closely with CD4 Th1 cells than with other CD8 T cell subsets. These results suggest that T cell states are more strongly shaped by functional status and signals from their microenvironment than by their lineage history.

Interestingly, progenitor CD8 T cells were most similar to CD4 Tfh cells in both their genomewide chromatin accessibility and transcriptomic profiles (**Fig. 3A, S8A,B**). These two subpopulations also exhibited shared activity (in both expression and chromatin accessibility) of key marker genes of CD8 progenitors and Tfh cells, including *Cxcr5*, *Bcl6*, *Tox*, *Pdcd1*, *Pou2f2*, *Tcf7*, and *Icos*, which distinguished them from other CD4 and CD8 T cell states (**Fig. 3B,C**).

On one hand, this similarity is consistent with the original characterization of progenitor exhausted CD8 T cells as a Cxcr5^+^ subpopulation within the CD8 lineage, while Cxcr5 is a canonical marker of CD4 Tfh cells, and with reports of other important molecules (e.g. Bcl6, Tcf1) shared between CD8 progenitors and Tfh cells.^8, 10, 41^ On the other hand, these two subpopulations have distinct functional roles shaped by their different lineages and immune contexts. Thus, this striking convergence between progenitor CD8 T cell and CD4 Tfh cell states motivated us to further investigate their shared regulatory programs at finer resolution.

### Archetypal analysis of scATAC-seq data reveals a mixed regulatory program of progenitor CD8 T cells

We sought to understand how transcriptionally and epigenomically similar phenotypes can arise in distinct T cell subsets, including across lineages. In particular, we wanted to further investigate the striking similarity between CD4 Tfh and progenitor CD8 T cell states. We hypothesized that these cell states converge on similar phenotypes through shared regulatory programs, driven by common TFs, cytokines, and signaling molecules. To characterize these shared molecular programs, we needed a framework beyond discrete, non-overlapping cell type annotations.

To achieve this, we turned to archetypal analysis of the scATAC-seq data (**Fig. 3D**). Archetypal analysis models chromatin accessibility profiles via a linear low-rank decomposition using a small number of archetypes, each representing an idealized regulatory program learned in an un-supervised manner from the data^53–55^ (**Methods**). The chromatin accessibility profile of each T cell can then be represented as a weighted combination of these archetypes. These archetypal weights provide a quantitative and interpretable framework for understanding how cell states relate to one another and how shared regulatory programs are deployed across cells. Each archetype is represented by a vector over accessibility peaks and can be interpreted post hoc based on the properties of the cells and peaks it comprises. We refer to the level of accessibility of a peak within an archetype as an archetypal score, reflecting the peak’s relative contribution to that archetypal regulatory program.

We applied archetypal analysis with *k* = 8 archetypes to our scATAC-seq data for T cells from the day 7 response to LCMV (**Fig. 3E-I, S8, S9**). Archetypes were annotated based on cell state annotations among cells with high archetypal weights, as well as functional annotations of genes whose expression was most strongly associated with the archetypes. Some archetypes corresponded predominantly to individual cell states, such as CD4 Treg or naive CD8 T cells, while others captured shared regulatory programs across functionally related subsets (**Fig. 3E,F**). For example, we identified a T cell activation archetype shared between CD4 Th1, CD8 effector and exhausted T cells, associated with high expression of proliferation genes *Top2a*, *Mki67* and *Birc5* and cytotoxic genes *Nkg7* and *Gzmb* (**Fig. 3E-G, S9, Methods**). We also distinguished between a CD8 effector archetype, predominant in effector CD8 T cells, and a separate CD8 hyperactivation or early exhaustion archetype (referred to as CD8 exhaustion), observed in exhausted CD8 T cells, but also in a subset of Th1 cells and in progenitor CD8 T cells. Notably, progenitor CD8 T cells lacked a dominant archetype and instead exhibited a mixed signature, primarily composed from the CD8 exhaustion and CD4 Tfh archetypes (**Fig. 3E,F, S8C,D**). This was consistent with their transcriptional and regulatory similarity to Tfh cells and their original definition as progenitor exhausted CD8 T cells. This pattern was robustly recapitulated when applying our archetypes to our reanalysis of the Daniel *et al.* and Giles *et al.* datasets^35, 36^ (**Fig. S8G,H, Methods**). Thus, archetypal analysis effectively captured the continuum of T cell phenotypes and revealed the hybrid regulatory program of progenitor CD8 T cells.

To better understand the regulatory content of each archetype, we examined archetypal scores at individual peaks (**Fig. 3D,H**). For interpretability, each peak was uniquely assigned to the archetype with the highest score for that peak (**Methods**). We then studied the relationship between scATAC-seq peak activity and scRNA-seq gene expression across cells. We found that for genes overexpressed in specific T cell subsets, nearby peaks assigned to the corresponding archetypes had significantly elevated accessibility (**Fig. 3I, S8F**). For example, peaks of the CD4 Tfh archetype were most accessible near genes highly expressed in CD4 Tfh cells, while peaks of CD8 effector and activation archetypes were enriched near genes upregulated in CD8 effector cells. Notably, genes overexpressed in progenitor CD8 T cells had equally strong association with both Tfh and exhausted CD8 T cell archetypes (**Fig. 3I**). Thus, the scATAC-seq peak archetypes captured biologically meaningful regulatory programs of gene expression regulation across T cell states and revealed mixed regulation of gene activity in progenitor CD8 T cells.

We next asked whether peak archetypes also reflect gene expression regulation *within* individual T cell populations. For many key T cell marker genes, we found that different chromatin accessibility peaks within the same gene locus were assigned to distinct archetypes (**Fig. 3H**), suggesting that these genes may be regulated by different sets of regulatory elements depending on cellular context. To explore this, we examined the correlation between the accessibility of individual peaks and gene expression within specific T cell subsets. For example, *Tox* is a gene expressed in CD4 Tfh and exhausted CD8 T cells, and is also highly expressed in progenitor Tcf1^+^ CD8 T cells^56–60^ (**Fig. 3B**). Among its 108 associated peaks, 49 were assigned to the CD4 Tfh archetype and 19 to the CD8 exhaustion archetype (**Fig. 3H, S10**). Within CD4 Tfh cells, *Tox* peaks assigned to the Tfh archetype were most frequently significantly correlated with *Tox* expression, whereas in exhausted CD8 T cells, peaks from the CD8 T cell exhaustion archetype showed the strongest correlation (**Fig. S10**). Thus, distinct regulatory elements, grouped by scATAC-seq archetypes, can be differentially associated with regulation of the same gene in a cell type-specific manner. This supports the view that the regulatory program of progenitor CD8 T cells integrates features of both CD4 Tfh and exhausted CD8 T cells, as reflected for example in the chromatin accessibility landscape of the *Tox* locus.

In conclusion, archetypal analysis of scATAC-seq data uncovered regulatory programs shared across T cell populations. These programs are closely linked to gene expression patterns, with the same gene regulated by distinct sets of peaks, assigned to different archetypes, in different T cell states. Archetypal analysis reveals that progenitor CD8 T cells exhibit a mixed regulatory program combining features of exhausted CD8 T cells and CD4 Tfh cells.

### TF motif analysis implicates regulatory drivers of archetypal programs and progenitor CD8 T cell identity

To better understand the transcriptional regulators shaping the landscape of continuous and discrete CD4 and CD8 T cell phenotypes during viral response, we next asked which transcription factors (TFs) are associated with the distinct chromatin accessibility archetypes described above, particularly those underlying the regulatory program of progenitor CD8 T cells.

To this end, we first identified putative TF targets by scanning genomic sequences within chromatin accessibility peaks for TF motif matches. We then used these motif-based TF target signatures to score individual cells in the scATAC-seq data and compared motif scores across T cell functional states and archetypes (**Methods**). To link TF activity with regulatory programs, we computed correlations between motif scores and archetypal weights across cells. Using this approach, we found that TF motifs most strongly enriched in progenitor CD8 T cells reflected a combination of those associated with the Tfh and CD8 T cell exhaustion archetypes (**Fig. 4A-E, S11A**). For example, progenitor cells were enriched with Nfat, Lef1/Tcf1, Pou2f1/2 (Oct1/2) motifs, associated with the CD4 Tfh archetype, and with Runx and Tbet (Eomes/Tbx21) motifs, associated with the exhaustion archetype. In addition, progenitor cells were also enriched with the AP-1 family motif, associated with both Tfh and exhaustion archetypes. These results further support the idea that progenitor CD8 T cells integrate regulatory programs from both Tfh cells and exhausted CD8 T cells.

**Fig. 4.**
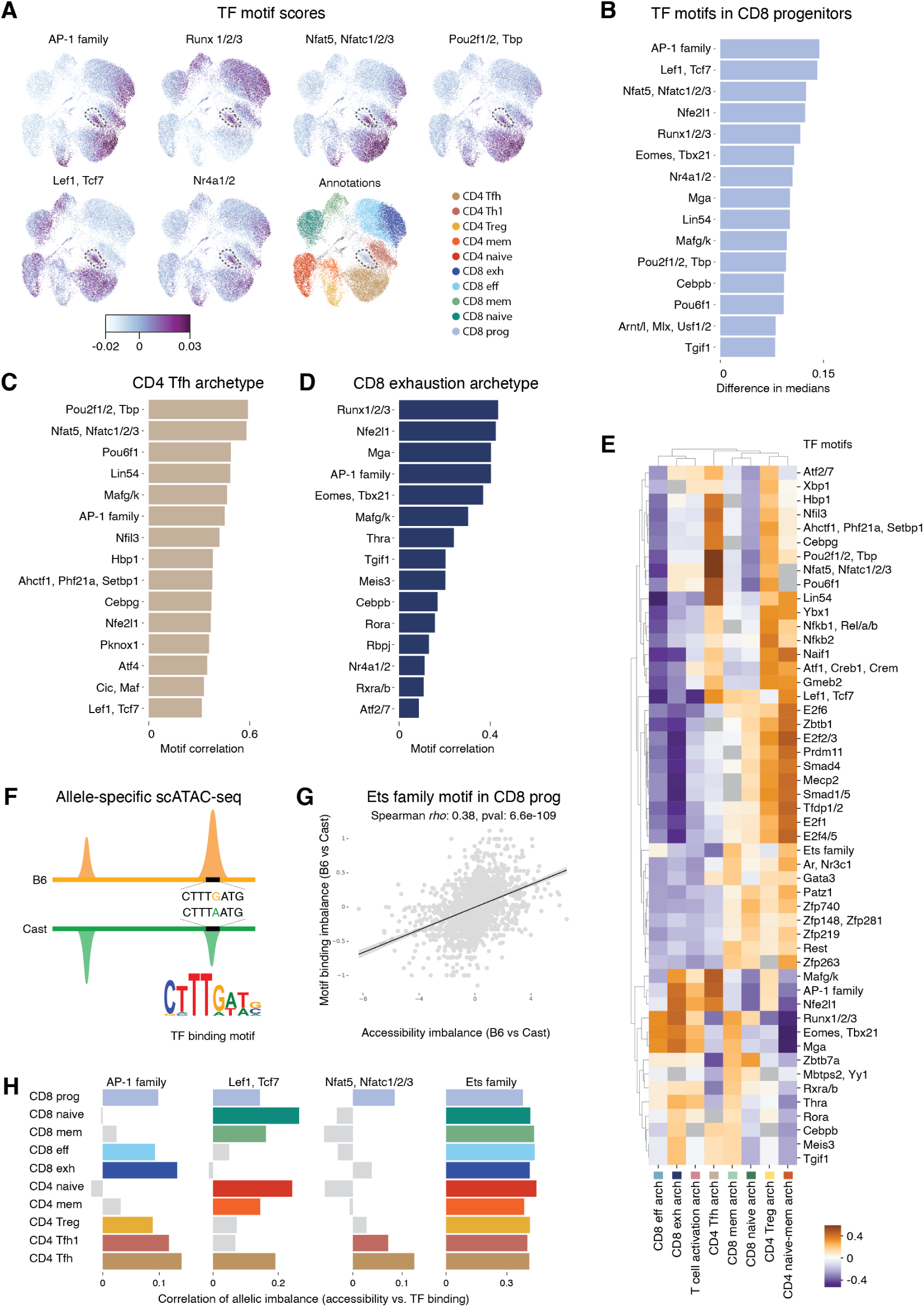
Distinct transcription factor motifs are associated with archetypal regulatory programs. **(A)** UMAP showing single-cell motif scores for selected transcription factor (TF) motifs (Methods). Cell type annotations are shown for reference. **(B)** Top 15 motifs significantly enriched in progenitor CD8 T cells compared to all other cells (FDR *≤* 0.05). **(C)** Spearman correlations between CD4 Tfh archetypal weights and motif scores across cells. Barplot shows the top 15 motifs with the strongest positive correlations. **(D)** Same as panel C, but for the CD8 T cell exhaustion archetype. **(E)** Clustermap of correlations between archetypal weights and motif scores (top 10 most correlated motifs per archetype). **(F)** Schematic of the allele-specific analysis used to associate TF motifs with chromatin accessibility in ATAC-seq peaks. Using sequence differences between B6 and Cast genomes, we assigned scATAC-seq reads to parental alleles and quantified allele-specific pseudobulk chromatin accessibility per cell type. In parallel, we assessed allele-specific differences in motif matches due to genetic variants between the two genomes. **(G)** Scatterplot showing allelic imbalance of chromatin accessibility (*x*-axis) vs. allelic imbalance of Ets motif matches (*y*-axis) across peaks in progenitor CD8 T cells. A significant positive correlation indicates that sequence variants with increased predicted Ets motif strength are associated with increased chromatin accessibility on the same allele, consistent with functional TF binding. **(H)** Spearman correlations between allelic imbalance of motif matches and chromatin accessibility across functional T cell states for selected motifs. Correlations that are not statistically significant (FDR > 0.05) are shown in gray.

To further validate the associations between TF motifs and T cell functional phenotypes, we turned to allele-specific analysis. Because our single-cell data were generated from F1 hybrid mice (B6 × Cast), we were able to assign scATAC-seq reads to parental alleles and quantify allele-specific chromatin accessibility at regulatory regions within each T cell state. In parallel, we used known sequence variants between the two genomes to compute allele-specific motif-based predictions of TF binding (**Fig. 4F,G**). We reasoned that if a TF is functionally active in a given cell population, the predicted allelic imbalance in its binding motif strength should correlate with observed allelic imbalance in chromatin accessibility.^44, 61^ Indeed, we observed a strong correlation for the Ets family motif across all T cell states (**Fig. 4G,H, S11C**), consistent with the widespread role of Ets family TFs in T cell regulation, modulated by co-factors in a cell type-specific manner.^44^ Whereas the above allele-agnostic motif enrichment analysis was performed across cell states (**Fig. 4A-E, S11B**), this allele-specific analysis, performed *within* each cell state, provided orthogonal support for motif activity patterns. Specifically, in progenitor CD8 T cells, we observed significant allele-specific correlations with accessibility for motifs such as Lef1/Tcf7, Nfat, and AP-1, indicating their functional engagement in this population (**Fig. 4H, S11C**). Comparing these allele-specific correlations across T cell subsets revealed that AP-1 motif was active in both Tfh and exhausted CD8 T cells, while Lef1/Tcf7 and Nfat motifs were selectively active in Tfh but not in exhausted CD8 T cells. This pattern reinforces the notion that progenitor CD8 T cells integrate regulatory programs from both Tfh and exhausted states. These findings motivated us to further investigate the extent of underexplored regulatory similarity between Tfh cells and progenitor CD8 T cells.

### Tfh archetype captures a similar gradient of differentiation within Tfh cells and progenitor CD8 T cells

We have shown that progenitor CD8 T cells share epigenomic and transcriptomic features with CD4 Tfh cells. To further investigate this connection, we focused on examining the epigenomic CD4 Tfh archetype and its relationship to the progenitor CD8 T cell population.

We first examined the behavior of the CD4 Tfh archetype within the CD4 Tfh cell population itself. Within this subset, we observed a continuous gradient of the Tfh archetypal weight (**Fig. 5A**). This gradient was positively correlated with expression of Tfh activation-associated genes such as *Tox*, *Pdcd1*, *Bcl6* and *Icos* and negatively correlated with quiescence-associated genes such as *Sell* and *Il7r* (**Fig. 5B,C**). This suggested that the Tfh archetype provides a quantitative measure of Tfh cell differentiation, or the strength of the Tfh phenotype, within the CD4 Tfh population. Furthermore, we observed significant positive and negative correlations between Tfh archetypal weight and the expression of numerous functionally important T cell genes (**Fig. 5C, S12A**). These included T cell receptor signaling genes *Zap70* and *Ptpn11*, co-stimulatory receptors *Icos*, *Tnfrsf4* and *Tnfsf8*, inhibitory molecules *Pdcd1*, *Tnfaip8* and *Nt5e*, and transcription factors *Bcl6*, *Tox*, *Hif1a*, *Ikzf2*, *Pou2f2*, *Id3*, *Nfatc1*, *Nfat5* and *Batf*. Consistently, the Tfh archetype was also correlated with the key TF motifs including Nfat, Lef1/Tcf1 and AP-1 family motifs (**Fig. 5D, S12F**). These associations further define the functional spectrum and regulatory drivers of the Tfh archetype and support its interpretation as a measure of progressive TCR-stimulated Tfh cell activation, differentiation and migration, enabling B cell help, while inhibiting proliferative expansion typical of effector cells.^10^ These results illustrate that scATAC-seq archetypal analysis can effectively capture a continuum of differentiation within the CD4 Tfh cell population.

**Fig. 5.**
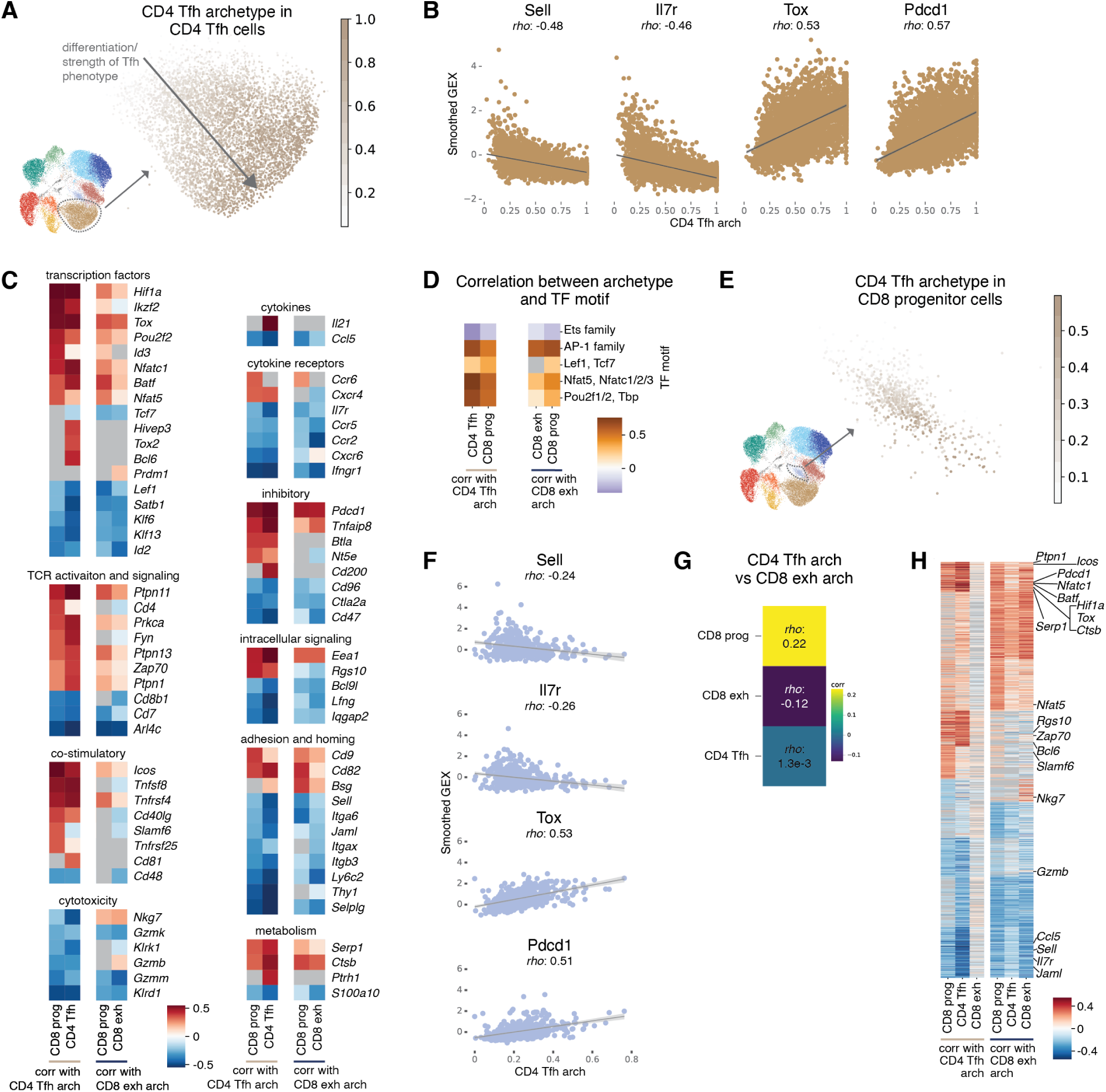
Tfh archetypal weight reflects similar gradient of differentiation in CD4 Tfh cells and progenitor Tcf1^+^ CD8 T cells. **(A)** UMAP showing CD4 Tfh archetypal weights in the CD4 Tfh cell population (excluding 61 outlier cells for visualization purposes). We propose that this Tfh archetypal gradient reflects Tfh differentiation and/or the strength of the Tfh phenotype. **(B)** Scatterplots of normalized expression of selected genes in CD4 Tfh cells along the Tfh archetypal gradient. The gene expression values were averaged over neighbors in the scRNA-seq *k*-nearest-neighbor (kNN) graph (*k* = 20). A smoothed fit using ordinary least squares is shown in gray. Spearman’s *ρ* shown on top. **(C)** Heatmap showing Spearman correlations between archetypal weights and normalized gene expression within selected cell populations for selected genes from different functional categories. Left two columns: correlations with the Tfh archetypal weights for progenitor CD8 T cells and CD4 Tfh cells. Right two columns: correlations with CD8 exhaustion archetypal weights for progenitor and exhausted CD8 T cells. The gene expression values were averaged over neighbors in the scRNA-seq kNN graph (*k* = 20). Non-significant correlations (FDR > 0.05) are shown in gray. **(D)** Heatmap showing Spearman correlations between archetypal weights and TF motif scores for selected motifs. Same archetypes and cell states as in panel C. Non-significant correlations (FDR > 0.05) are shown in gray. **(E)** UMAP showing the Tfh archetypal weights in the progenitor Tcf1^+^ CD8 T cell population (excluding 9 outlier cells for visualization purposes). **(F)** Scatterplots of normalized expression of selected genes in progenitor CD8 T cells along the Tfh archetypal gradient. The gene expression values were averaged over neighbors in the scRNA-seq kNN graph (*k* = 20). A smoothed fit using ordinary least squares is shown in gray. Spearman’s *ρ* shown on top. **(G)** Heatmap comparing the Spearman correlations between Tfh and CD8 exhaustion archetypal weights calculated within three cell populations. **(H)** Heatmap showing Spearman correlations between either Tfh or CD8 exhaustion archetypal weights and normalized gene expression, within CD4 Tfh, progenitor, and exhausted CD8 T cells. Genes with the strongest positive or negative correlations between exhausted CD8 T cells and CD8 exhaustion archetype, CD4 Tfh cells and Tfh archetype, and CD8 progenitors and either archetype were selected (total 1,131 genes). The gene expression values were averaged over neighbors in the scRNA-seq kNN graph (*k* = 20).

Given the prominent contribution of the CD4 Tfh archetype to the regulatory phenotype of progenitor CD8 T cells (**Fig. 3E,F, 4B,C**), next we wanted to quantitatively explore the Tfh archetype within the progenitor population. Strikingly, we found that the progenitor CD8 T cells exhibited a continuous gradient of Tfh archetypal weights, similar to that observed within CD4 Tfh cells (**Fig. 5C-F, S12A,C,F**). In particular, many genes showed comparable correlations with the Tfh archetype in both CD4 Tfh population and the CD8 progenitor population, reinforcing the regulatory similarity between the two populations.

We also examined the contribution of the CD8 exhaustion archetype to the progenitor population. The exhaustion archetype was uncorrelated with the Tfh archetype both across the full dataset and within individual cell states including Tfh cells, exhausted CD8 T cells and progenitor CD8 T cells (**Fig. 5G, S12E**). Some key T cell genes, most notably the Tfh- but also exhaustion-associated genes *Tox*and *Pdcd1*, showed similarly high expression correlations with both archetypes, particularly within progenitor cells (**Fig. 5C,H**). This was consistent with the functional role of progenitors as activated cells that must balance responsiveness with restraint of proliferation. Nevertheless, the majority of genes most correlated with the Tfh and CD8 exhaustion archetypes were distinct (**Fig. 5H, S12F**), indicating that the two archetypes capture largely non-overlapping regulatory programs.

Together, these findings suggest that the progenitor CD8 T cell state reflects a unique integration of regulatory inputs from both Tfh- and exhaustion-like programs. Using scATAC-seq archetypal analysis, we reveal a Tfh-like gradient of differentiation within progenitor CD8 T cells, supporting the view that these cells, similar to CD4 Tfh cells, adopt an activated identity while avoiding full effector-like differentiation and proliferation.

### Clone 13 infection accelerates progression along the Tfh differentiation gradient in CD4 Tfh and progenitor CD8 T cells

While we have consistently observed similar T cell states and phenotypes in both LCMV Arm and Cl13 infections, the immune responses to these clones ultimately diverge over time. To investigate the basis of these differences, we examined how epigenomic and regulatory programs vary between the two infections. We focused particularly on the shared population of progenitor CD8 T cells, as they can give rise to both effector-like and terminally exhausted CD8 T cells and are therefore central to understanding how T cell response evolves in each context.

We found that within the progenitor CD8 T cell population, the Tfh and CD8 exhaustion archetypes were significantly higher in the Cl13 response compared to Arm, whereas the CD8 naive archetype was significantly higher in Arm (**Fig. 6A,B, S8C**). Similar trends—stronger activation in Clone 13 and a more naive-like state in Armstrong—were also observed in other antigen-experienced T cell populations (**Fig. 6A,B, S8C**). This was consistent with prior reports that the overall T cell response to Cl13 is more strongly activated than to Arm and ultimately progresses towards exhaustion as a protective mechanism to limit immunopathology.^7, 8^

**Fig. 6.**
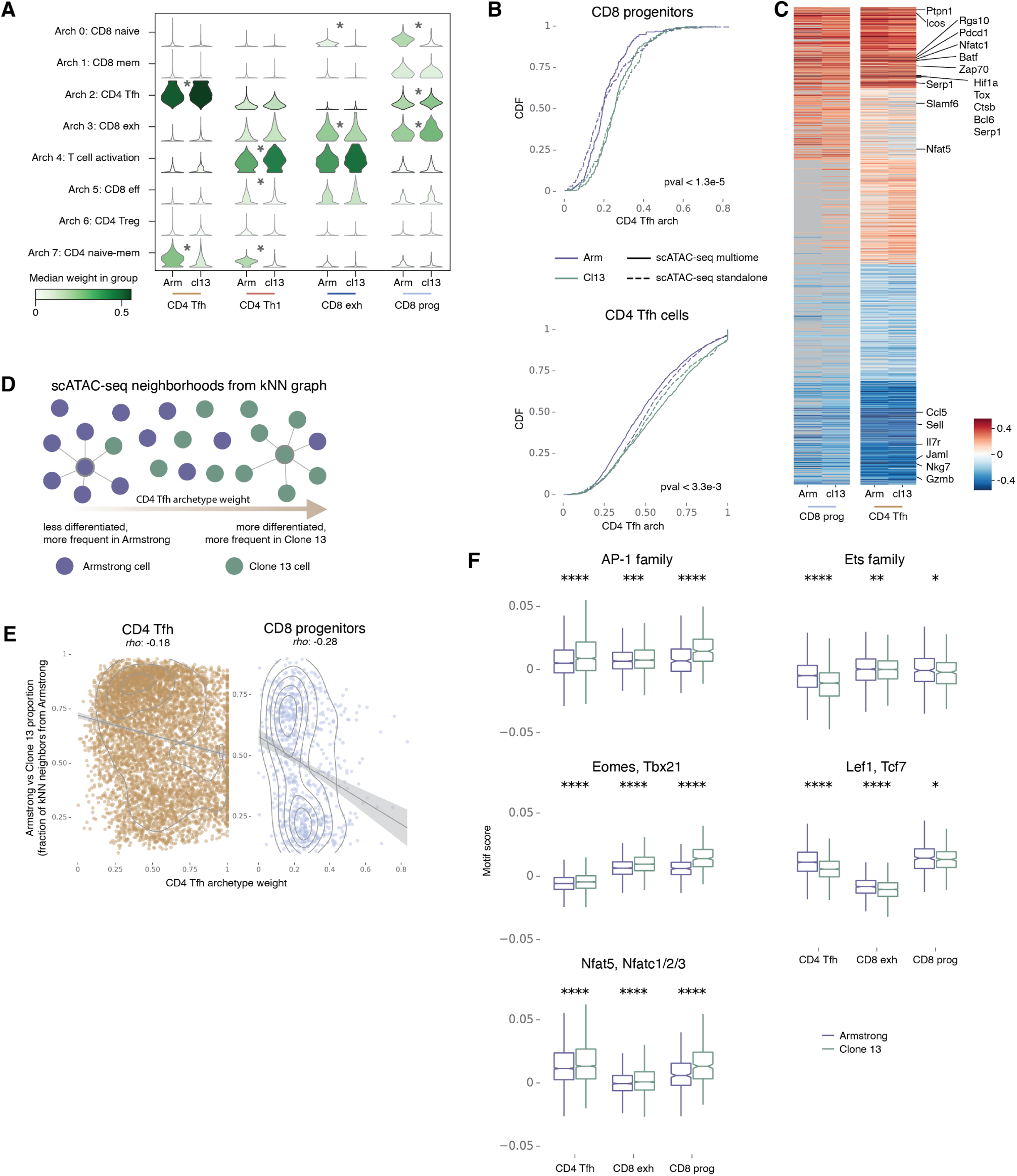
Clone 13 infection accelerates progression along the Tfh differentiation gradient in CD4 Tfh and progenitor CD8 T cells. **(A)** Violin plots showing the distributions of archetypal weights for selected T cell subsets, separately in Arm and Cl13 infections. Significant differences between infections within a population are marked with asterisks (FDR *≤* 0.05, KS test; absolute median difference *≥* 0.05). **(B)** CDF curves showing the distribution of the Tfh archetypal weights for Arm and Cl13 cells in progenitor CD8 T cell population (top) and the CD4 Tfh cell population (bottom), separately for standalone and multiome scATAC-seq data. Twosided KS test p-value listed (after Benjamini-Hochberg correction) for all pairwise comparisons across all batches between Armstrong and Clone 13 cells. **(C)** Heatmap showing the correlations between the Tfh archetypal weights and normalized gene expression in progenitor CD8 T cells and CD4 Tfh cells, separately for Arm and Cl13 infections, for the same genes as in Fig. 5H. **(D)** Schematic of the comparison of cells along the Tfh archetypal gradient between Arm and Cl13 infections. For each cell, the fraction of neighbors (i.e. most similar cells by scATAC-seq profile) from each infection is computed based on the scATAC-seq kNN graph constructed for all samples. We propose that cells from Clone 13 infection progress more rapidly along the Tfh differentiation trajectory as captured by the Tfh archetypal weights. **(E)** Scatterplot of the Tfh archetypal weights (*x* axis) and the Arm vs. Cl13 proportions (*y* axis) in CD4 Tfh cells (*left* ) and progenitor CD8 T cells (*right* ). The Arm vs. Cl13 proportion is calculated as the fraction of cell neighbors in the scATAC-seq kNN graph (*k* = 100) that belong to the Armstrong samples, and then smoothed over 30 nearest scATAC-seq neighbors. **(F)** Distribution of selected TF motif scores for different T cell populations, colored by infection (Arm vs. Cl13). Asterisks in the plot represent statistical significance (two-sided KS test, after BH correction) of the difference between the two infection types within a population (*: FDR *<* 0.2, **: FDR *<* 0.05, ***: FDR *<* 0.01, ****: FDR *<* 0.001). Boxplots: center line, median; box limits, upper and lower quartiles; whiskers, 1.5*×* interquartile range; outliers removed for clarity.

We then examined more closely the Tfh-like differentiation trajectory within progenitor CD8 T cells that we identified (**Fig. 5**). A similar Tfh differentiation gradient was observed in both Arm and Cl13 responses, within both CD4 Tfh and progenitor CD8 T cells, as evidenced by comparable gene expression correlations (**Fig. 6C, S12A**). However, consistent with the overall stronger T cell activation observed in Cl13, cells from the Cl13 response progressed more rapidly along this gradient than those from Arm (**Fig. 6B,D,E**). This finding was robust across the two scATAC-seq profiling technologies (standalone and multiome). TF motif analysis implicated potential regulators of these differential responses between Arm and Cl13. In particular, activationassociated motifs AP-1, Tbet and Nfat showed stronger signal in Cl13, while motifs associated with naive and memory-like states, such as Lef1/Tcf7 and Ets motifs, were more enriched in Arm (**Fig. 6F, S13E**). These observations were confirmed in reanalysis of published scATAC-seq datasets (**Fig. S13A-D**). Thus, we conclude that although progenitor CD8 T cells exhibit a similar Tfh-like differentiation gradient in both Arm and Cl13 responses, they advance along this gradient more rapidly in Cl13 infection. This suggests a potential for further divergence of the T cell response upon persistent antigen stimulation.

## Discussion

In this study, we integrated single-cell chromatin accessibility and transcriptomic profiling with a novel comprehensive reference of T cell regulatory regions to analyze CD4 and CD8 T cell responses to acute and chronic viral infection. Our approach enabled high-resolution, interpretable identification of all major T cell states, including naive, memory, effector, Tfh, regulatory, exhausted, and progenitor populations. We observed that each of these populations was present across both infection types, but at differing frequencies.

We focused in particular on the clinically relevant progenitor Tcf1^+^ CD8 T cell population. While the relationship between progenitor populations in acute and chronic infection has been a key area of investigation,^19, 32, 35–40^ our analysis demonstrated that these cells form a highly similar population at the transcriptomic and epigenomic levels between the two infection settings.

Surprisingly, we found that these CD8 progenitors exhibit striking regulatory similarity with CD4 Tfh T cells, a population with a key role in B cell help via direct physical interactions in the B cell follicles, and defined by sustained TCR signaling and restrained effector differentiation. Despite arising from the distinct lineages of CD4 and CD8 T cells, both populations share chromatin accessibility and transcriptomic profiles and co-activate transcription factors such as Tcf1, Bcl6, Nfat family factors and Tox.

To mechanistically dissect this convergence, we applied archetypal analysis to the chromatin accessibility data, revealing that progenitor CD8 T cells do not represent a unique regulatory program, but rather a composite of CD8 exhaustion and Tfh archetypes. This supports a modular view of progenitor identity, arising from reuse of regulatory programs deployed in other T cell states. Concordantly, via TF motif analysis, we observed that progenitor CD8 T cells combine activity of Tfh-enriched TFs including Lef1, Tcf1, Pou2f2 and Nfat family, exhaustion-associated factors with Runx and Tbet motifs, and AP-1 family factors with activity in both CD8 exhaustion and Tfh cells. This decomposition analysis suggests that Nfat family factors, triggering Tox and Pd1 activation, may be involved in functionality and adaptations characteristic of both Tfh cells and CD8 progenitors, such as restrained effector differentiation immediately upon and later during persistent TCR stimulation. Activity of other factors, including AP-1 family factors e.g. Batf, may be associated with functional activation program shared by all antigen-experienced CD4 and CD8 T cells that is progressively lost during differentiation towards exhaustion; Tbet and Runx factors may be associated with the preparation for and execution of cytotoxic effector functions. This view is consistent with and helps further refine known functionality of these factors in T cells,^19, 62–68^ though more specific functional studies of these and other factors are needed to further dissect combinatorial mechanistic regulatory programs across T cell populations.

These findings suggest possible functional and spatial links between progenitor CD8 T cells and Tfh cells. Future studies should explore whether they co-localize and interact, and thus converge to activation of similar regulatory programs despite originating from different T cell lineages. Consistently, a similar subset of CD8 T cells expressing CXCR5, exhausted genes, and memory-like genes have been discovered to localize in the B cell follicles, i.e. along with Tfh cells, but were originally defined as a less-exhausted, follicular CD8 T cell population.^69–73^ Our results suggest that progenitor and follicular CD8 T cells may be the same or similar populations. Indeed, CD4 T cells and B cells influence the progenitor CD8 T cells during infection, though it remains unclear whether CD4 T cell help directly affects the formation or maintenance of the CD8 progenitors.^24, 74–77^ However, progenitor CD8 T cells were reported to localize in T cell zones, away from B cell follicles, at later stages of chronic infection, suggesting potential heterogeneity within the progenitor population or a dependence of spatial localization on the stage of infection progression.^41, 67, 78, 79^ Further research using multi-omics, spatial transcriptomics and proteomics, and direct cell-cell interaction measurements such as uLIPSTIC,^80^ across time courses and various systems of T cell stimulation and response, could help better characterize the T cell phenotypic landscape and explore the localization and interactions of Tfh cells and progenitor and follicular CD8 cells. Perturbation experiments, including targeted depletion of one population or blockade of shared signaling pathways or transcriptional regulators could further shed light on their relationship.

In sum, our work highlights the power of multiomic and computational approaches to dissect complex T cell regulation and offers a generalizable framework for studying shared and distinct programs across immune contexts. Applying this strategy to other T cell settings, such as tumorinfiltrating lymphocytes, vaccine responses, or autoimmune diseases, could uncover novel regulatory states and reveal common principles of immune cell plasticity. These approaches will also be broadly applicable across a range of biological contexts beyond immunology.

## Methods

### Single-cell multiomics

Experimental procedures for mouse work, cell isolation and single-cell genomics were as previously described.^44^ All animal experiments in this study were approved by the Sloan Kettering Institute (SKI) Institutional Animal Care and Use Committee under protocol 08-10-023. Specifically, Foxp3^GFP-DTR^ B6/Cast F1 males were infected with LCMV Armstrong (2 × 10^5^ pfu I.P.) or LCMV Clone 13 (2 × 10^6^ pfu I.V./retroorbital). Spleens from 3 mice (Armstrong group) or 4 mice (Clone 13 group) were pooled. T cells were enriched using anti-CD4 and anti-CD8 dynabead flowcomp kits (mixing CD4 and CD8 antibody). Following enrichment, cells were stained with the following panel of antibodies and BV510 ghost dye: NK1.1: BV421; CD62L: BV605; CD4: Percp-Cy5.5; CD44: APC; TCRb: PE-Tx-red. A 50-50% mix of CD4 and CD8+ T cells (2 million cells total) was sorted into complete RPMI with 10% FCS, washed once, resuspended at 1M cells/ml and used as input material for single cell analysis.

For the standalone scATAC-seq samples, libraries were prepared according to the 10x scATAC-seq protocol from 10x Genomics Chromium (Single Cell ATAC Reagent Kits User Guide (catalog no. CG000168, Rev A)). Briefly, cells were centrifuged (300g, 5min, 4°C) and permeabilized with 100*µ*l of chilled lysis buffer (10mM Tris-HCl, pH 7.4, 10mM NaCl, 3mM MgCl2, 0.1% Tween-20, 0.1% IGEPAL CA-630, 0.01% digitonin and 1% bovine serum albumin). The sample was incubated on ice for 3–5min and resuspended with 1ml of chilled wash buffer (10mM Tris-HCl, pH 7.4, 10mM NaCl, 3mM MgCl2, 0.1% Tween-20 and 1% bovine serum albumin). After centrifugation (500g, 5min, 4°C), the pellet was resuspended in 100 *µ*l of chilled Nuclei Buffer (catalog no. 2000153, 10x Genomics). Nuclei were counted using a hemocytometer, and finally the nucleus concentration was adjusted to 3,000 nuclei per *µ*l. We used 15,360 nuclei as input for tagmentation. Nuclei were diluted to 5 *µ*l with 1× Nuclei Buffer (10x Genomics) and mixed with ATAC buffer (10x Genomics) and ATAC enzyme (10x Genomics) for tagmentation (60 min, 37 °C). ScATAC-seq libraries were generated using the Chromium Chip E Single Cell ATAC kit (10x Genomics, catalog no. 1000086) and indices (Chromium i7 Multiplex Kit N, Set A, 10x Genomics, catalog no. 1000084) following the manufacturer’s instructions. Final libraries were quantified using a Qubit fluorimeter (Life Technologies) and the nucleosomal pattern was verified using a BioAnalyzer (Agilent). Libraries were sequenced on a NovaSeq6000 (Illumina) with the following read lengths: 50+8+16+50 (Read 1+Index 1+Index 2+Read 2).

For the multiomic scATAC-seq + scRNA-seq samples, libraries were prepared according to the Chromium Next GEM Single Cell Multiome ATAC + Gene Expression User Guide (CG000338 Rev A). Briefly, cells were centrifuged (300g, 5 min, 4°C) and permeabilized with 100 *µ*l chilled lysis buffer (10mM Tris-HCl pH 7.4, 10mM NaCl, 3mM MgCl2, 0.1% Tween-20, 0.1% IGEPAL-CA630, 0.01% digitonin, 1% BSA). The samples were incubated on ice for 3-5 min and resuspended with 1 ml chilled wash buffer (10mM Tris-HCl pH 7.4, 10mM NaCl, 3mM MgCl2, 0.1% Tween-20 and 1% BSA and RNase Inhibitor 1 U/µL; Lucigen NxGen, F83923-1). After centrifugation (500g, 5 min, 4°C), the pellets were resuspended in 100 *µ*l chilled Nuclei buffer (2000153, 10x Genomics). Nuclei were counted using a haemocytometer, and finally the nucleus concentration was adjusted to 3,000 nuclei per *µ*l. 15,400 permeabilized nuclei were diluted to 5 *µ*L in isotonic Tagmentation Buffer (20 mM Tris-HCl pH 7.4, 150 mM NaCl, 3 mM MgCl2, RNase Inhibitor 1 U/*µ*L; Lucigen NxGen, F83923-1). Nuclei were mixed with 10 *µ*L of a transposition master mix consisting of 7 *µ*L ATAC Buffer B (10x Genomics, 2000193) and 3 *µ*L ATAC Enzyme B (Tn5 transposase; 10x Genomics, 2000265) per reaction. Transposition was performed at 37°C for 60 min, followed by a brief hold at 4°C. Multiomic libraries were generated using the Chromium NextGEM Chip J (10x Genomics, 2000264).

For read alignment and initial generation of count matrices, cellranger-atac and cellranger-arc count were run twice for each sample, once with the standard 10x mm10 genome and a second time with a custom Cast genome reference to enable allele-specific analysis. For the scRNA-seq modality, a final allele-specific BAM file for each sample was generated by combining the B6 and Cast GEX BAM file output by the two runs of CellRanger. In this combined BAM file, each read was labeled either with the genome that had the higher mapping alignment score or as ambiguous if the mapping scores were similar. Similarly for the scATAC-seq data, an allele-specific BAM file was generated for each sample by combining the B6 and Cast ATAC BAM file output by the two runs of CellRanger, labeling reads as B6, Cast, or ambiguous based on alignment scores. A final BED file of single nucleotide insertion sites (shifted by +4/-5) was generated from each BAM file.

### ATAC-seq peak atlas for T cells

#### Bulk ATAC-seq peaks

We built our T cell peak atlas by extending a previously published compendium of CD8 T cell peaks.^32^ We generated a new compendium of CD4 T cell peaks using seven previously published CD4 T cell studies (**Table S1**). For these studies, FASTQ files were downloaded from SRA. Bowtie2^81^ was used to align the FASTQ files for each sample to mm10 genome using the flags --no-unal, --no-mixed, and --no-discordant. Samtools^82^ was used to turn the SAM files produced by Bowtie2 into sorted BAM files. The sample BAM files were then turned into BED files using BEDtools^83^ bamtobed function, and then shifted by +4/-5. For samples with technical replicates, the shifted BED files were concatenated together and sorted. MACS2^84^ was then run on the shifted BED files for each sample, using the flags --nomodel, --shift 0, --extsize 76, -p 0.1, -B, --SPMR, and --keep-dup ‘auto’. In addition, MACS2 was run on all of the shifted BED files for all samples in a study, using the same flags. To identify peaks that were reproducible across biological replicates, IDR v2.0.3^85^ was then run using all biological replicates of a sample from a study (denoted with the --samples flag) versus the background peak list run on all samples in the study (denoted with the --peak-list flag). The final IDR peaks for each study were then sorted and merged together using bedtools.

#### Single-cell ATAC-seq peaks

An initial analysis of the scATAC-seq component of our multiomic dataset was performed using ArchR^86^ on the standard mm10 CellRanger output in order to filter out low-quality cells and generate a single-cell peak set. An ArchR project was generated using ArrowFiles with all default parameters. A tile matrix was generated using tileSize of 500 and binarize set to False. Cells were filtered out based on low number of nFrags, with different thresholds used for each of the four samples (1200 for multiome Armstrong, 1500 for multiome Clone 13, 5000 for scATAC Armstrong, and 6000 for scATAC Clone 13). Doublets were called using the addDoubletScores function with k = 15. Cells with a DoubletEnrichment score greater than or equal to 3 were filtered out, removing 3,819 cells. Iterative LSI was performed with the addIterativeLSI function, called with iterations = 8, clusterParams = list(resolution = c(1.5), sampleCells = 10000, n.start = 10, maxClusters = 25), totalFeatures = 1*e*08, varFeatures = 5*e*06, force = TRUE, and firstSelection = “Var”. Batch correction was performed using the addHarmony function^87^ in ArchR on the IterativeLSI dimension-reduced matrix, for the four samples (scATAC Armstrong, scATAC Clone 13, multiome Armstrong, multiome Clone 13). A UMAP was generated on the batch-corrected Harmony reduced dimensions, using the addUMAP function. A gene score matrix was generated with the addGeneScoreMatrix function and all default parameters, and clusters were generated using the addClusters function with resolution = 1.2 and reducedDims=“Harmony”. Clusters were assigned to different T cell states based on marker gene scores from the GeneScoreMatrix, and one cluster was found to be B cells.

Peaks on the single-cell data were then called using MACS2 for all T cells using a BED file of single base pair insertion sites with the flags --nomodel, --shift -75, --extsize 150, -q 0.001, -B, --SPMR, and --keep-dup ‘all’.

#### Merging bulk and single-cell ATAC-seq peaks

The high quality bulk ATAC-seq peaks for CD4 T cells from IDR analysis were read in to R as GRanges^88^ objects and combined using the GenomicRanges::reduce function, to generate a simplified set. These were then filtered to peaks with a size greater than 76.

This peak set, a set of previously published peaks from CD8 T cell data,^32^ and the scATAC-seq peaks were combined using the GenomicRanges::reduce function, and again filtered to peaks with a size greater than 76. This generated a set of 223,820 peaks, which was then filtered to only those on the standard chromosomes, resulting in a final set of 223,492 peaks (**Table S2**).

Nearest genes to peaks were determined with a custom script using the bedtools^83^ nearest function.

#### Peak signatures

We learned bulk ATAC-seq peak signatures by using DESeq2^89^ to compare chromatin accessibility between all samples from pairs of conditions within a study. For each study, peaks were filtered to those where at least 20% of the samples had counts greater than or equal to 10 reads. Then, differentially accessible peaks were calculated using DESeq2 between the two conditions. Significantly differentially accessible peaks between the two conditions of interest in either direction (log2 fold change *>* 1, adjusted p-value ≤ 0.05) were stored as epigenomic peak signatures (**Table S3**). In the scATAC-seq data, these peak signatures were scored using score_genes function in scanpy.^90^

### scATAC-seq data analysis

Peak counts were generated separately for each of the four samples (standalone scATAC-seq Armstrong, multiome scATAC-seq Armstrong, standalone scATAC-seq Clone 13, and multiome scATAC-seq Clone 13) in an allele-specific way to count reads aligned to the B6 genome separately from reads aligned to the Cast genome, using a custom script from the single basepair BED file of Tn5 insertions. The allele-specific counts were then summed together for each sample to generate total counts (across both alleles). The scanpy^90^ package was used for all downstream analyses. Cells were filtered out if they had low total counts across all genes, or low number of peaks with non-zero count per cell. Peaks were filtered so that they were present in at least 0.5% of cells in any one of the four samples, resulting in a dataset of 200,771 peaks and 32,100 cells. Fragment counts were generated by taking the ceiling of peak counts / 2, to remove the bias of even-numbered counts.^91^ Pearson residual normalized values^92^ were then generated separately for each of the four samples, using theta parameter for overdispersion set to 0.3 for the two scATAC-seq standalone samples and 1 for the two scATAC-seq multiome samples (theta was chosen such that the distributions for normalized peak means was centered on 0 and normalized peak variances were centered around 1). Two additional peaks were filtered out due to non-finite values after Pearson residual normalization, resulting in a dataset of 200,769 peaks. Then PCA was run with 100 PCs using the Pearson residual values, and due to a batch effect between multiome vs. standalone scATAC-seq, Harmony^87^ was run to integrate the multiome and standalone components of the data. A kNN graph of single-cell chromatin accessibility similarities was built on the Harmony-integrated PCA matrix using 30 PCs, 30 nearest neighbors and the cosine similarity metric. Leiden clustering was performed with a resolution of 0.7, and UMAP was generated with the default parameters. One cluster (cluster 11) was removed due to high accessibility at peaks near B cell genes, suggesting this cluster was likely B cell contamination, resulting in a dataset of 31,888 cells. A new neighborhood graph was built using 30 nearest neighbors, 30 PCs, cosine metric on the Harmony-integrated PCA matrix, Leiden clustering was rerun with resolution of 1, and UMAP was recalculated with default parameters. An additional cluster (cluster 15) was filtered out due to high doublet scores from running the scrublet^93^ function in scanpy. This resulted in a final dataset of 31,662 cells and 200,769 peaks. Cells were assigned to annotations based on accessibility at marker genes and using the epigenomic signature scores from our bulk ATAC-seq compendium. Specifically, to distinguish CD4 and CD8 T cell lineages, we used both gene expression and chromatin accessibility of the lineage-defining genes *Cd4* and *Cd8a*. We identified quiescent naive and memory CD4 and CD8 T cells, likely not actively responding to LCMV, based on activity of *Sell*, *Lef1*, *Tcf7*, *Slamf6* and *Ifngr2*. Among CD4 T cells, we defined regulatory T cells (Treg) using *Foxp3*, follicular helper T cells (Tfh) using *Bcl6*, *Cxcr5*, *Pdcd1*, *Tox* and *Tcf7*, and activated Th1 cells using *Ifng*. Among CD8 T cells, we distinguished between progenitor cells associated with *Cxcr5*, *Slamf6*, *Tcf7*, *Pdcd1* and *Tox*, and a continuum of actively responding cells. We classified them into hyperactivated cells in the early state of exhaustion (referred to as exhausted from now on), more frequent in clone 13 response and associated with higher activity of inhibitory and exhaustion marker genes *Pdcd1*, *Tox*, *Havcr2* (*Tim3* ), *Lag3* and *Tigit* and elevated cytotoxicity as indicated by overexpression of *Gzmb*, *Nkg7* and *Prf1*, and the remaining effector cells more frequent in the Armstrong response. Enrichment of bulk ATAC-seq peak signatures in these scATAC-seq clusters further confirmed these functional annotations.

### ATAC-seq peak summits and TF binding analysis

For each of the four scATAC-seq samples, the 1 bp-resolution Tn5 insertion BED file was intersected with the peak regions using bedtools intersect. These were then sorted, and a bedGraph file was generated from the BED using a custom script. The bedGraph file was then turned into a bigWig using the bedGraphToBigWig^94^ method. The bigWig files were read into R as RleLists, the signal was accumulated over the four samples, and then subset to the peak regions. Using the biosignals^95^ package, 150 bp windows of maximum signal within the peaks were uncovered. These summit regions were then used to predict TF binding to both the B6 and Cast genomes using the package motifmatchr^96^ and a p-value threshold of 1e−3. For allele-specific analysis, imbalanced TF motif match was determined as the difference in predicted binding between B6 and Cast, with an absolute (non-zero) difference less than 4 (i.e. the sites with the absolute difference *>* 4 were excluded from the analysis as outliers). For allele-agnostic TF motif analysis, the maximum predicted binding value to either the B6 or Cast genome was used for a TF motif in a summit. To explore these TF motifs in our scATAC-seq data, we first looked for enrichment of TF motifs in differentially accessible peaks for each cell type (pval ≤ 0.05, LFC *>* 0). We used the hypergeometric test to determine enrichment of summits within differentially accessible peaks with predicted TF motif matches (as in **Fig. 4H, S11F**). In addition, cells were scored for TF motifs by using scanpy score_genes function with the set of peaks that contain at least one summit with a TF motif match (see **Fig. 4A**).

### Allele-specific analysis

Allele-specific counts were generated separately for the four samples over 475,357 summit windows within peaks. Counts were either assigned to the B6 genome, the Cast genome, or were considered ambiguous and ignored in downstream analyses. The scanpy package was used for all downstream analyses, and an AnnData object was generated by concatenating all four samples and then filtered to cells in our final dataset. Counts were then normalized by first generating a scaling factor for each cell, summing read counts for B6 and Cast counts. The scaling factor for each cell was then divided by the mean scaling factor over all cells. Finally the B6 and Cast count matrices were separately divided by the cell-based scaling factor. Allele-specific imbalance values (log2 fold changes) were then generated using these normalized counts to compare accessibility for B6 reads vs. Cast reads within each cell state. P-values were generated using a paired Wilcoxon test, and then corrected using Benjamini-Hochberg multiple hypothesis correction, and considered significantly imbalanced if FDR ≤ 0.05. We used Spearman correlation to compare allelic imbalance between predicted TF motif matches and chromatin accessibility. Correlations in each cell state were calculated only for motifs with at least 1000 imbalanced summits.

### scRNA-seq data analysis

The cell by gene count matrices were generated using a custom script, using only exonic reads. All analyses were performed using scanpy. QC steps were performed separately for the Armstrong samples and Clone 13 samples. Cells were first filtered out if having high mitochondrial UMI counts (using standard CellRanger filtered matrix output). Then cells were also filtered out if containing a low number of genes with non-zero count (fewer than 200), and genes were filtered out if present in fewer than 3 cells in either sample, resulting in 15,397 genes for further analysis. Cells were filtered out if having large total counts (25,000 for the clone 13 sample and 35,000 for the Armstrong sample). Pearson residual normalization was performed on each sample separately with theta = 10 for both. The two samples were then concatenated together, resulting in a final dataset of 19,186 cells. Standard single-cell RNA-seq analysis steps were run: PCA, kNN neighborhood graph construction with 100 neighbors and 40 PCs and cosine similarity metric, Leiden clustering with resolution of 1.5, and UMAP generated. One cluster was filtered out as likely B cell contamination (high *Cd79a* expression), resulting in 19,107 cells. Cells were then filtered out if not present in the scATAC-seq data component, and cell type annotations from the scATAC-seq analysis were used to annotate cells. This resulted in a final dataset of 18,972 cells. Cell cycle gene signature scores were generated using scanpy score_genes_cell_cycle function and the G2M and S genes from cc.genes lists in the package Seurat.^97^

### Archetypal analysis

Archetypal analysis of the scATAC-seq data was run using the py_pcha package^54^ using the transposed Pearson residual normalized count matrix. We used *k* = 8, 12, and 15, and chose *k* = 8 for further analysis. In order to characterize archetypal peaks, the matrix of archetype scores (*H* ∈ ℝ *^k^*^×^*^p^*, where *p* is the number of peaks) was used to uniquely assign peaks to the archetype with the highest value (if greater than the threshold value of 0.1). To apply archetypes to other datasets (such as the Daniel *et al.*^35^ and Giles *et al.*^36^ datasets, as in **Fig. S8G,H**), scanpy gene_score function was used to score the peaks assigned to each archetype (**Table S6**). These scores were then linearly scaled so that they were in the 0-1 range and summing to 1 for each cell. To generate gene expression vectors for each archetype, we calculated weighted averages of cell expression vectors. In archetypal analysis, we decompose our original cell by peak (*n* × *p*) data matrix into two matrices (see **Fig. 3**).^55^ More explicitly, *X* ≈ *WH*, where *X* is our original cell by peak (R*^n^*^×^*^p^*) normalized count matrix, *W* ∈ ℝ *^n^*^×^*^k^* is the archetypal weights (providing the probability each cell is assigned to each archetype), and *H* ∈ ℝ *^k^*^×^*^p^* is the archetypal score matrix, representing accessibility profiles for each archetype. The archetype scores matrix *H* is generated directly from the original data matrix: *H* = *BX*, because each archetype profile is a linear composition of the cells in the dataset. We used the weights from the *B* ∈ ℝ *^k^*^×^*^n^* matrix (representing how strong an effect each cell has on an archetype) to calculate the weighted average of expression profiles for each archetype, only for cells in the multiome.

For Gene Ontology enrichment analysis of genes highly correlated with the Tfh archetype (**Fig. S12B**), the GProfiler^98^ tool (using the python interface, v1.0.0) was run separately for the 100 top and bottom correlating genes, and run separately for each database source, with a significance threshold of 0.05. The background set of genes was all genes in our scRNA-seq dataset.

### Analysis of other datasets

**Daniel *et al.***^35^ Fragment files were downloaded from SRA for day 8 samples (chronic gp33+, chronic gp33-, acute gp33+, and acute gp33-). BED files with single base pair Tn5 insertion sites were generated from the fragment files using a custom script, and then intersected with our peak atlas BED file using bedtools intersect. Counts were generated over all peaks in our peak atlas using a custom script from the Tn5 insertion BED file. The QC analysis was done for each of the four samples separately, with cells being filtered based on the total read counts, and then peaks filtered based on presence in sufficient number of cells across all samples. Fragment counts were generated from raw counts using the suggested ceiling of peak counts / 2 to remove the bias towards even counts, and then normalized with Pearson residuals, as described above. The samples were then concatenated, PCA was run, kNN neighborhood graph was generated using 30 neighbors and 40 PCs with cosine similarity metric, and UMAP and Leiden clustering with resolution of 0.3 were run. Scoring of progenitor vs. exhaustion peak signatures was calculated using scanpy default score_genes function.

**Giles *et al*** ^36^ Fragment files were downloaded from SRA for three samples: Armstrong infection 8 days post infection, Clone 13 infection 8 days post infection, and naive cells. BED files with single base pair Tn5 insertion sites were generated from the fragment files using a custom script, and then intersected with our peak atlas BED file using bedtools intersect. Counts were generated over all peaks in our peak atlas using a custom script. Basic QC steps (filtering cells due to low total counts and peaks due to low counts across cells) were run in scanpy, separately for each of the three samples. Peak counts were first fragment-adjusted (ceiling of peak counts/2) to remove the bias towards even counts and then normalized with Pearson residuals, as described above. Samples were then concatenated, resulting in a matrix of 36,959 cells and 144,833 peaks. PCA was run, kNN graph was generated with 30 neighbors, 40 PCs, and cosine similarity metric, UMAP was run with default parameters, and Leiden clustering was run with a resolution value of 0.4. Scoring of progenitor vs. exhaustion peak signatures was calculated using scanpy default score_genes function.

## Acknowledgments

We thank all members of the Pritykin lab for helpful discussions. This work was supported by the NIH/NIAID grant DP2AI171161, AACR-Bristol-Myers Squibb Immuno-oncology Research Fellowship (19-40-15-PRIT) and the Ludwig Institute for Cancer Research (Y.P.); Boehringer Ingelheim, the Austrian Science Fund (FWF, 10.55776/PAT4163824), and the ERC grant ERC-2023-STG 101116251 (J.v.d.V); NIH grants P30 CA008748 and R01 AI034206 (A.Y.R.). A.Y.R. is an investigator with the Howard Hughes Medical Institute. We thank the Single Cell Analytics Innovation Lab at MSKCC and the head of the lab Ronan Chaligné for processing single cell multi-omics samples.

## Supplementary Figures

**Fig. S1.**
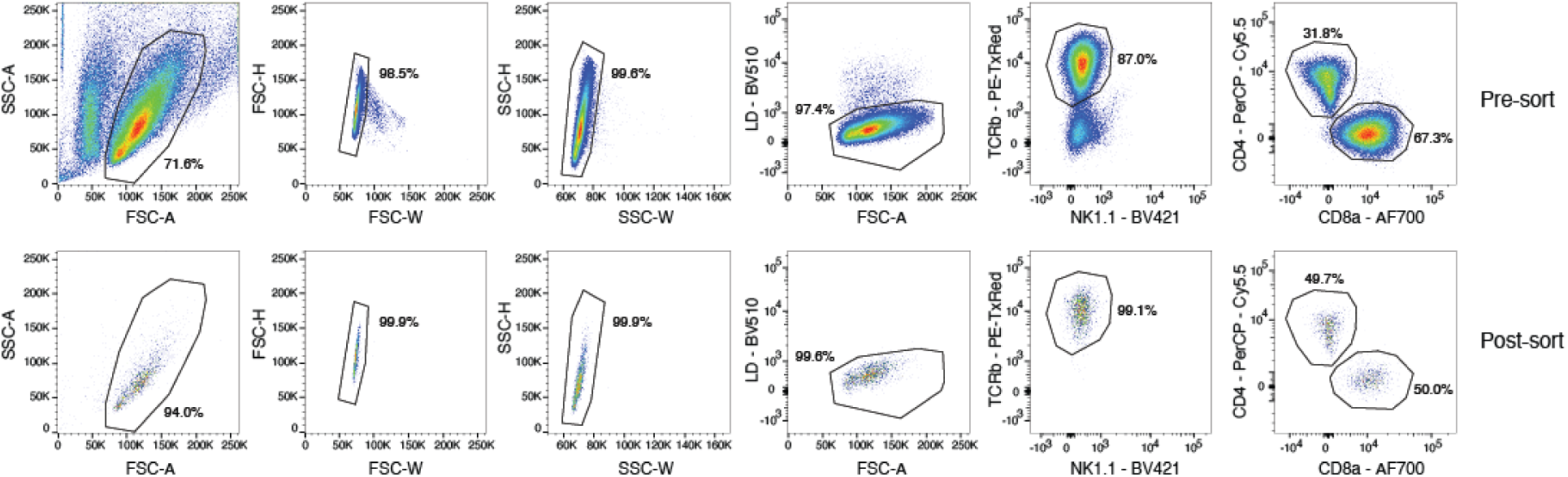
Flow cytometry sorting strategy (related to Figure 1). Splenic CD4 and CD8 T cells from mice infected with LCMV were isolated for multi-omic profiling (see **Methods**).

**Fig. S2.**
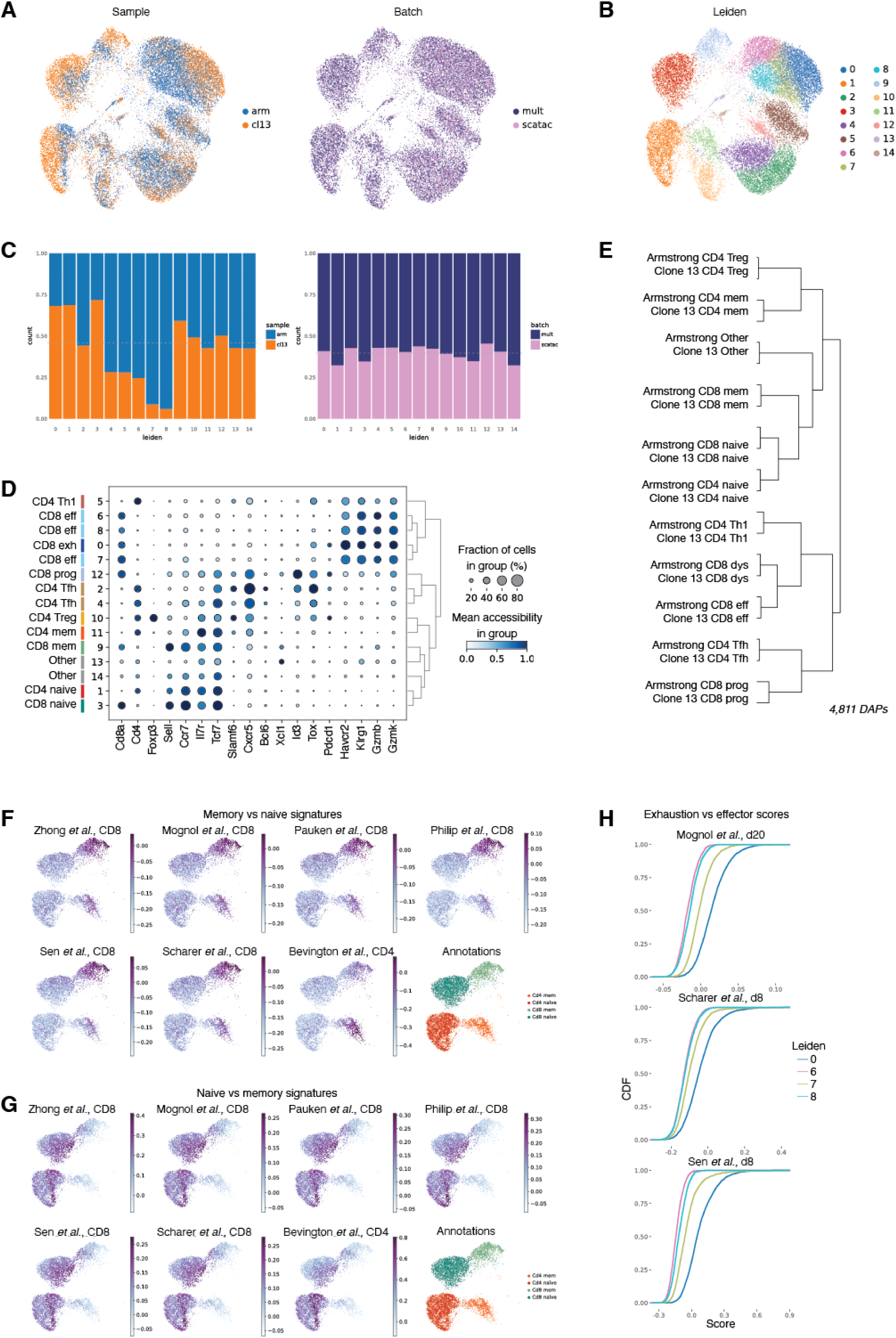
Analysis and functional annotation of the scATAC-seq data (related to Figure 1). **(A)** UMAP plot of the scATAC-seq component of the data, colored by (*left* ) sample (infection type) and (*right* ) experimental batch (scATAC-seq from multiome and standalone scATAC-seq experiment). **(B)** UMAP colored by Leiden clusters. **(C)** Bar plots showing composition of Leiden clusters by (*left* ) infection type and (*right* ) batch (scATAC-seq from multiome vs. standalone scATAC-seq). **(D)** Dotplot showing normalized scATAC-seq accessibility of a selected peak for each marker gene across scATAC-seq Leiden clusters. Cluster assignments to functional cell states are shown on the left. A dendrogram on the right shows how Leiden clusters relate to each other. Size of the dot represents the fraction of cells with normalized accessibility greater than 0, and the color of the dot represents the mean accessibility for a peak in a cluster. **(E)** Dendrogram of hierarchical clustering over 4,811 differentially accessible peaks (peaks enriched in at least one cluster, with log2 fold change >1.5 and *p ≤* 0.05), for T cell functional states, separated by infection type. **(F-G)** UMAP of the four clusters annotated as naive and memory populations, colored by (F) memory vs. naive and (G) naive vs. memory epigenomic signatures derived from different bulk ATAC-seq dataset (Methods). Shown also is a UMAP with final functional cell state annotations. **(H)** Cumulative distribution function (CDF) curves showing the distribution of three exhausted vs. effector epigenomic signature scores derived from bulk ATAC-seq data, separately for each of the four clusters annotated as either effector or exhausted CD8 T cells (distribution of cluster 0 is significantly different from all other clusters for all scores, FDR < 2.4e*−*112).

**Fig. S3.**
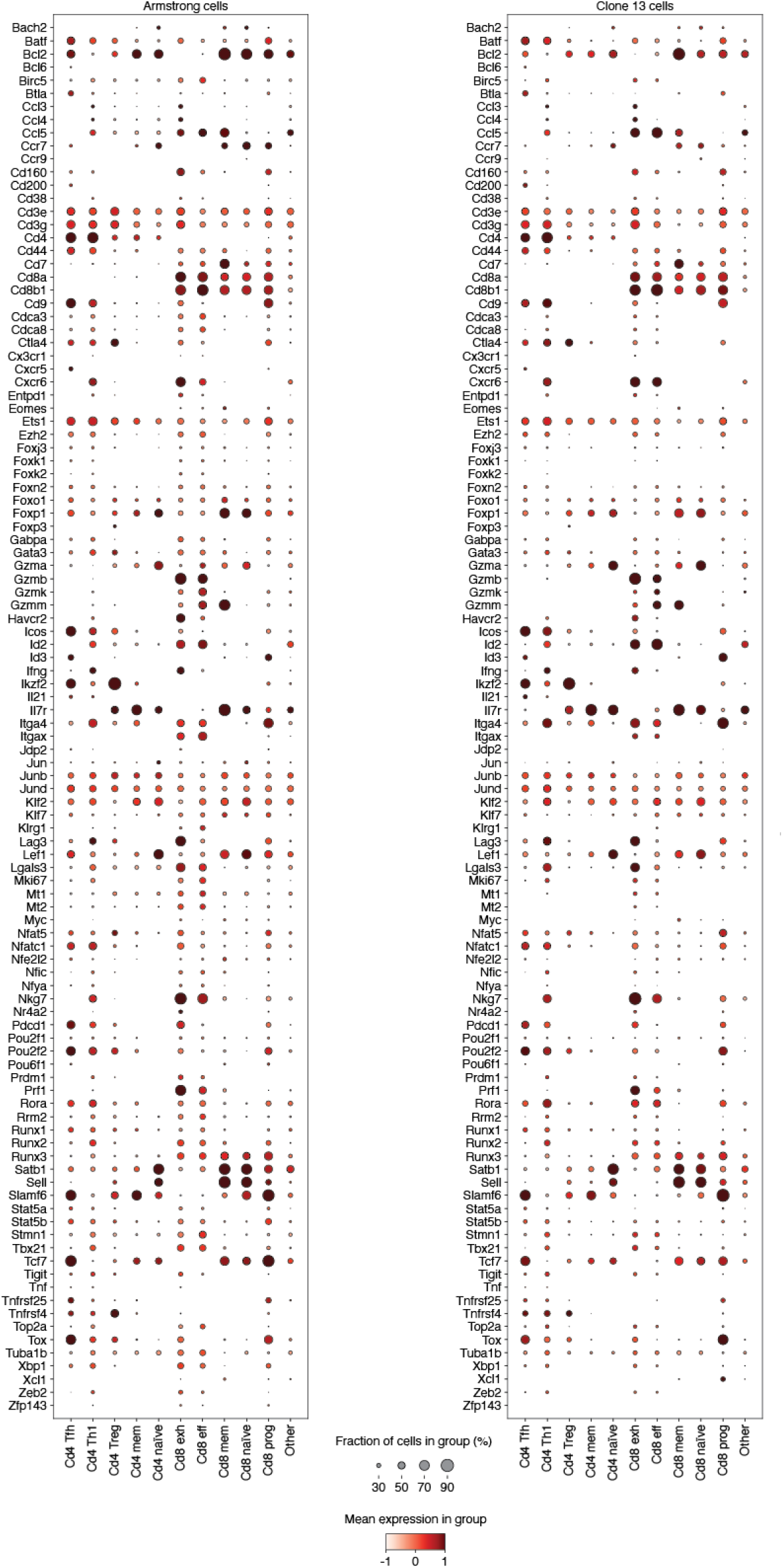
Expression of selected genes across functional cell states (related to Figure 1). Dotplot showing normalized scRNA-seq expression of genes with important functions in T cells, across functional cell states derived from scATAC-seq analysis, separated by infection type (left, Arm; right, Cl13). Dot size indicates the fraction of cells with normalized gene expression greater than zero; color represents the mean gene expression within each group.

**Fig. S4.**
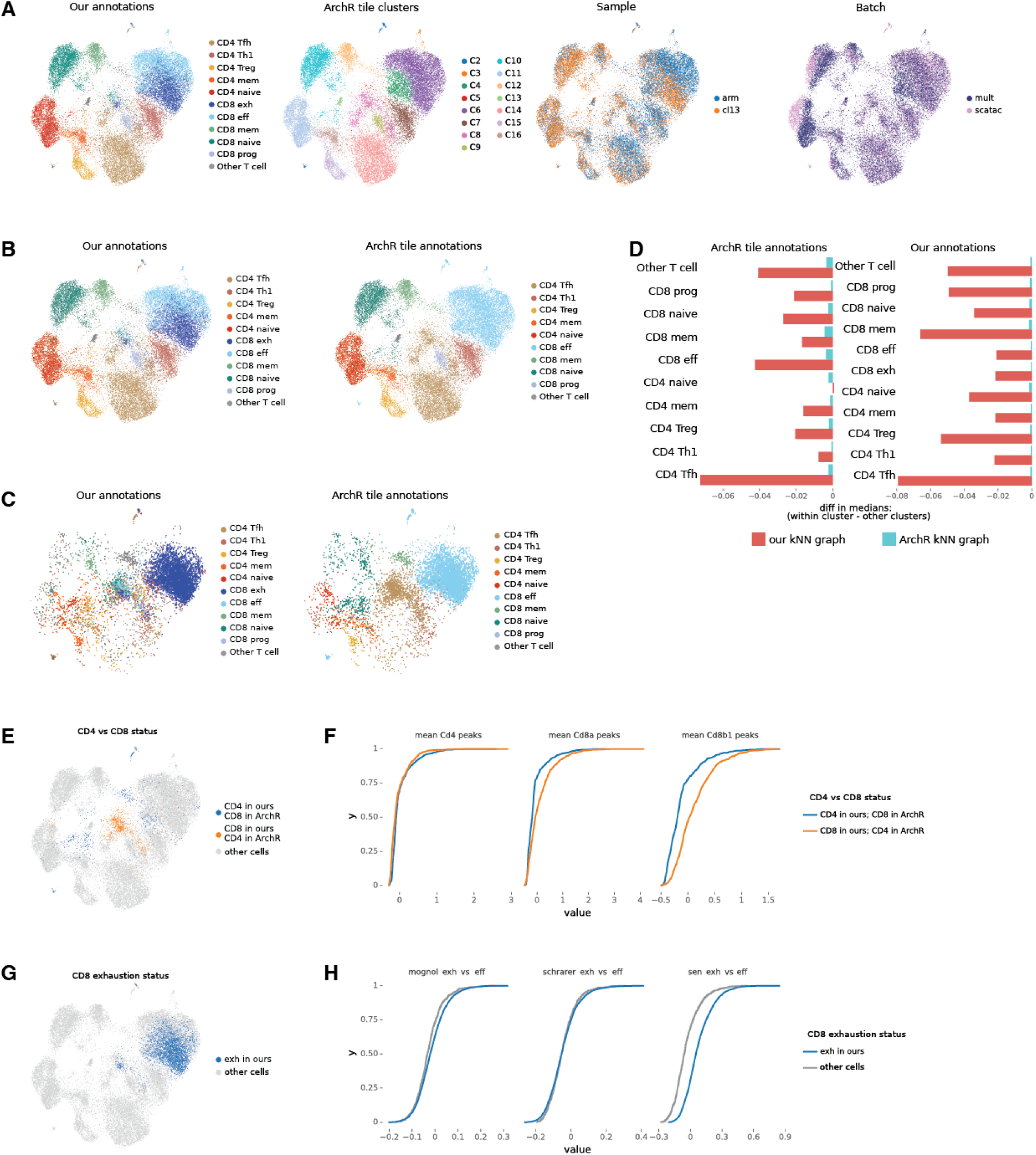
Comparison of our scATAC-seq analysis with ArchR tile-based analysis (related to Figure 1). **(A)** UMAP generated using the default ArchR pipeline with a TileMatrix for downstream analyses. UMAP plots are colored by (*left to right* ): our peak-based annotations, ArchR tile-based clusters, sample (infection type), and experimental batch (multiome scATAC-seq vs. standalone scATAC-seq). **(B)** ArchR UMAP colored by our cell state annotations and by annotations derived from ArchR tile-based clustering. ArchR clusters were annotated based on predominant overlapping annotations from our analysis. **(C)** ArchR UMAP plots showing cells with conflicting annotations between our analysis and the ArchR tile-based analysis. **(D)** Comparison of scATAC-seq similarities for cells sharing the same annotation versus cells with different annotations. scATAC-seq kNN graphs were built using low-dimensional representations from either ArchR tile-based LSI or our PCA analysis (30 dimensions, 20 nearest neighbors). For a kNN graph, for each cell from a certain functional cell state, we calculated the average distance to its neighbors from the same cell state and the average distance to its neighbors from other cell states, and then took the difference in medians for these two values (Methods). We calculated this both using ArchR based annotations (*left* ) and our annotations (*right* ). A negative value indicates higher similarity between cells with the same annotations. We found a significant difference in distances to neighbors from the same vs. different cell state between our kNN graph and the ArchR kNN graph for all annotations, FDR < 0.005. **(E, F)** Comparisons of cells that are annotated as CD4 vs. CD8 T cells in our vs. ArchR annotations (1,514 cells). **(E)** UMAP plots. **(F)** CDF plots showing distributions of accessibility for cells from each annotation group, for peaks of the lineage-defining genes *Cd4*, *Cd8a*, and *Cd8b1* (all differences are significant, FDR < 5.4e*−*19, two-sided KS test). Accessibility values were averaged across peaks near each gene for each cell. **(G, H)** Comparison of cells that are annotated as exhausted CD8 T cells in our analysis but not in ArchR annotations, as there were no clusters with this annotation in the ArchR analysis (4,075 cells). **(G)** UMAP plots. **(H)** CDF plots showing distributions of exhaustion epigenomic signature scores (derived from our bulk ATAC-seq compendium) for cells annotated as exhausted CD8 T cells in our analysis versus those annotated as effector CD8 T cells in both analyses (gray line). Blue values are significantly larger than gray values for all three scores (FDR < 3.1e*−*58, two-sided KS test).

**Fig. S5.**
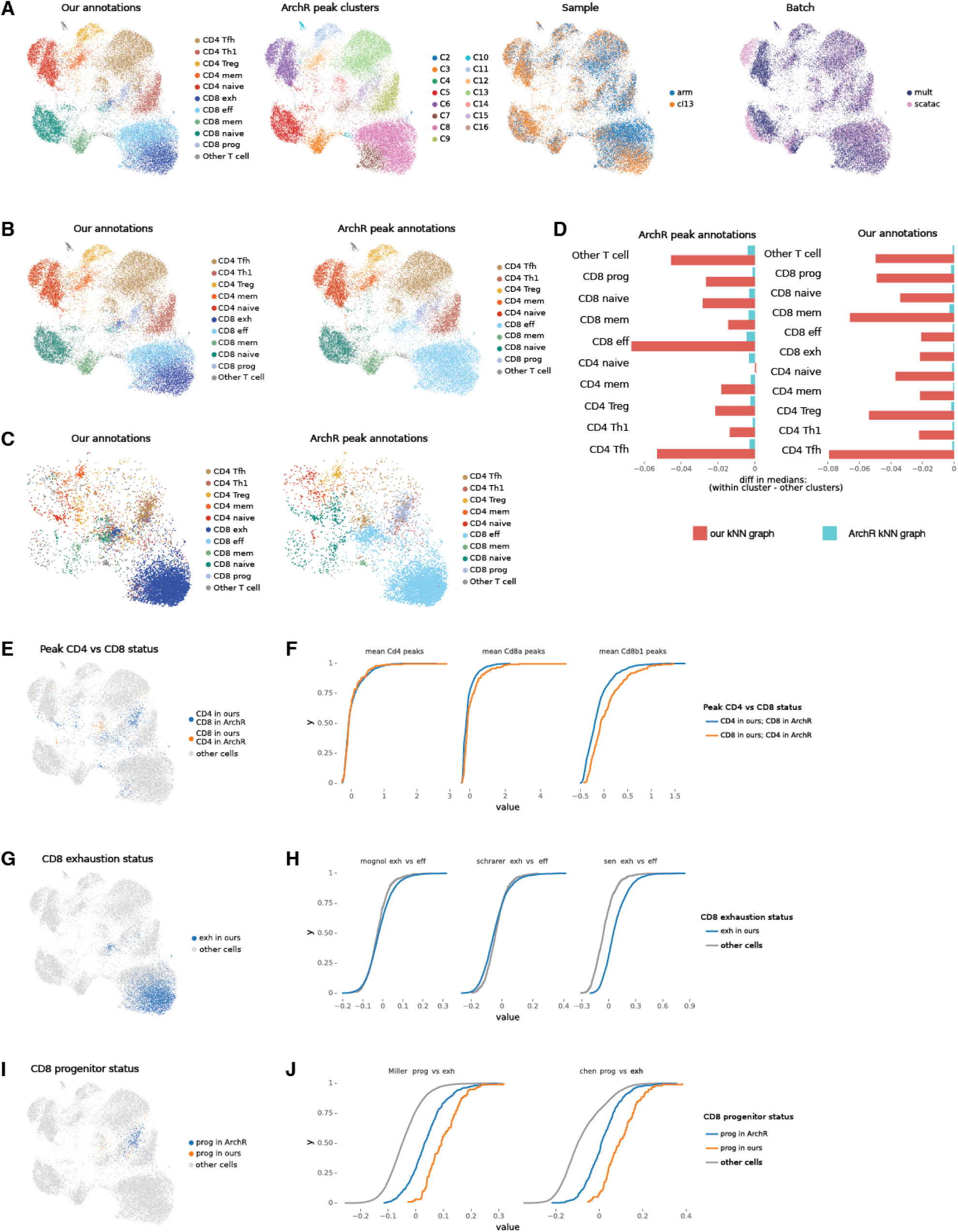
Comparison of our scATAC-seq analysis with ArchR peak-based analysis (related to Figure 1). Same comparative analysis as in Fig. S4 but for the peak-based instead of tile-based ArchR analysis. **(A)** UMAP generated using the default ArchR pipeline using an ArchR-generated PeakMatrix with MACS2 peak calling. UMAP plots are colored by (left to right): our peak-based cell state annotations, ArchR peak-based clusters, sample (infection type), and experimental batch (multiome scATAC-seq vs. standalone scATAC-seq). **(B)** ArchR peak-based UMAP colored by our cell state annotations and by annotations derived from ArchR peak-based clustering. ArchR clusters were annotated based on predominant overlapping annotations from our analysis. **(C)** ArchR peak-based UMAP plots showing cells with conflicting annotations between our analysis and the ArchR peakbased analysis. **(D)** Comparison of neighborhood distances for cells sharing the same annotation versus cells with different annotations. Analogous to Fig. S4D, but comparing with ArchR peak-based annotations rather than tile-based. We found a significant difference in distances to neighbors from the same vs. different cell state between our kNN graph and the peak ArchR kNN graph for all annotations, FDR < 0.008. **(E, F)** Comparisons of cells that are annotated as CD4 vs. CD8 T cells in our vs. ArchR peak-based annotations (1,353 cells). Analogous to Fig. S4E,F. The two distributions are significantly different for Cd8a and Cd8b1 (FDR < 1.7e*−*10, two-sided KS test). **(G, H)** Comparisons of cells that are annotated as exhausted CD8 T cells in our analysis but not in ArchR peak-based annotations, as there were no clusters with this annotation in the ArchR peak-based analysis (4,075 cells). Analogous to Fig. S4G,H. Blue values are significantly larger than gray values for all comparisons (FDR < 2.7e*−*97, two-sided KS test). **(I, J)** Comparisons of cells that are annotated as progenitor CD8 T cells in either our analysis or ArchR peak-based analysis but not both (608 cells) **(I)** UMAP plots. **(J)** CDF plots showing distributions of progenitor vs. exhausted epigenomic signature scores (derived from our bulk ATAC-seq compendium) for cells with conflicting annotations between our and ArchR analysis. Cells are either annotated as progenitor CD8 T cells in our analysis and not in ArchR, as progenitor cells in ArchR analysis and not our analysis, or non-progenitors in both analyses (gray line). The orange values are significantly larger than blue values in both comparisons (FDR < 2.4e*−*12, two-sided KS test).

**Fig. S6.**
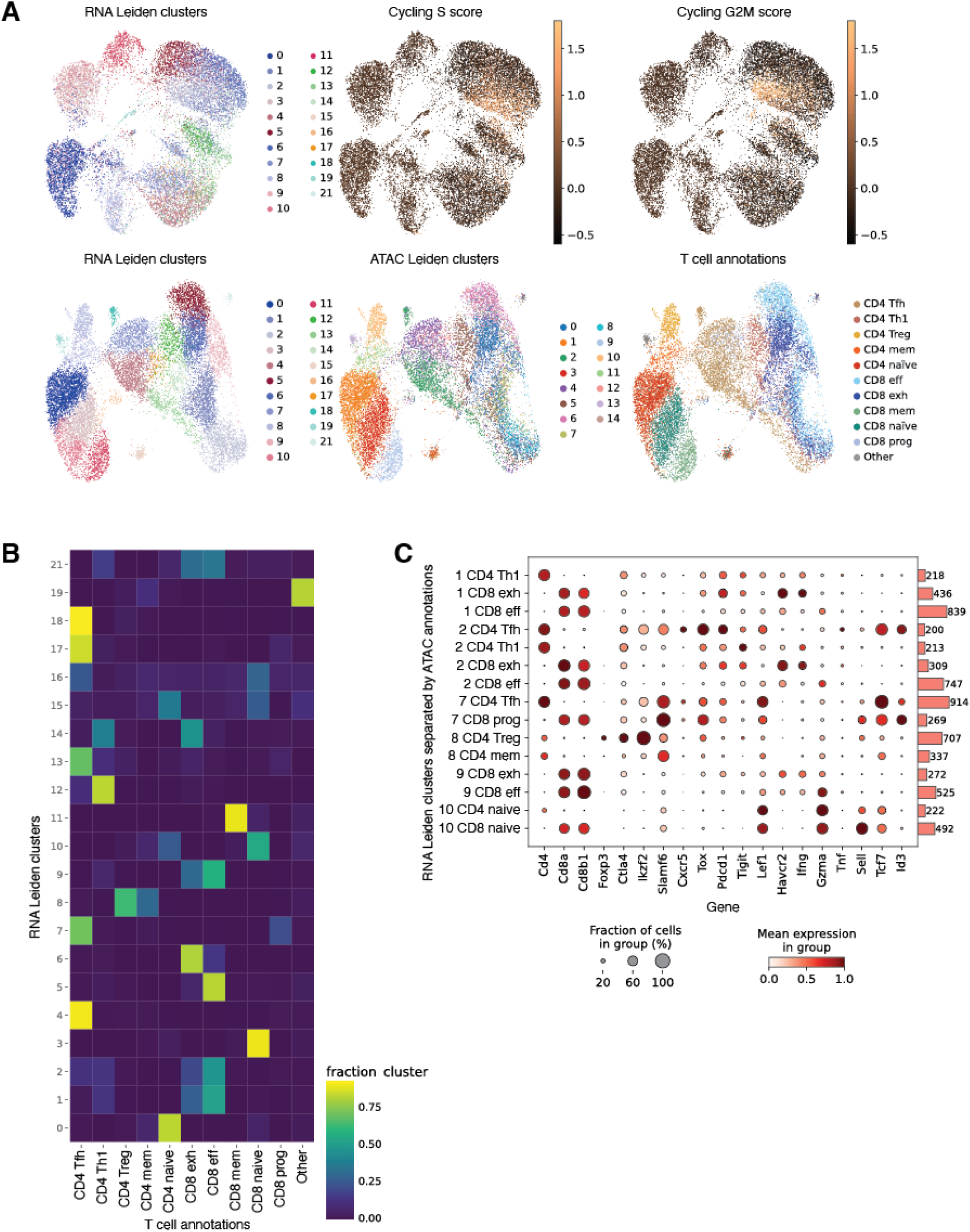
Analysis of the scRNA-seq data (related to Figure 1). **(A)** *Top row* : scATAC-seq UMAP plots colored by (*left* ) scRNA-seq Leiden clusters and (*center, right* ) cell cycle gene signature scores (S phase and G2M phase) in the scRNA-seq data. *Bottom row* : scRNA-seq UMAP plots colored by (*left* ) scRNA-seq Leiden clusters,(*center* ) scATAC-seq Leiden clusters and (*right)* our final functional T cell state annotations (based on the scATAC-seq analysis). **(B)** Heatmap showing the correspondence between scRNA-seq-based Leiden clusters and our final cell state annotations (based on the scATAC-seq analysis). Each square represents the fraction of cells from a given scRNA-seq cluster assigned to each annotation; rows sum to 1. **(C)** Dotplot showing expression of selected marker genes across scRNA-seq-based Leiden clusters, further stratified by their assigned scATAC-seq-derived functional annotations. Only groups of at least 200 cells are shown. Dot size indicates the fraction of cells with normalized gene expression greater than zero; color represents the mean gene expression within each group. The bar plot on the right shows the number of cells assigned to each scATAC-seq annotation. We conclude that scRNA-seq clustering cannot resolve between distinct functional annnotations, including between cells from CD4 and CD8 lineages.

**Fig. S7.**
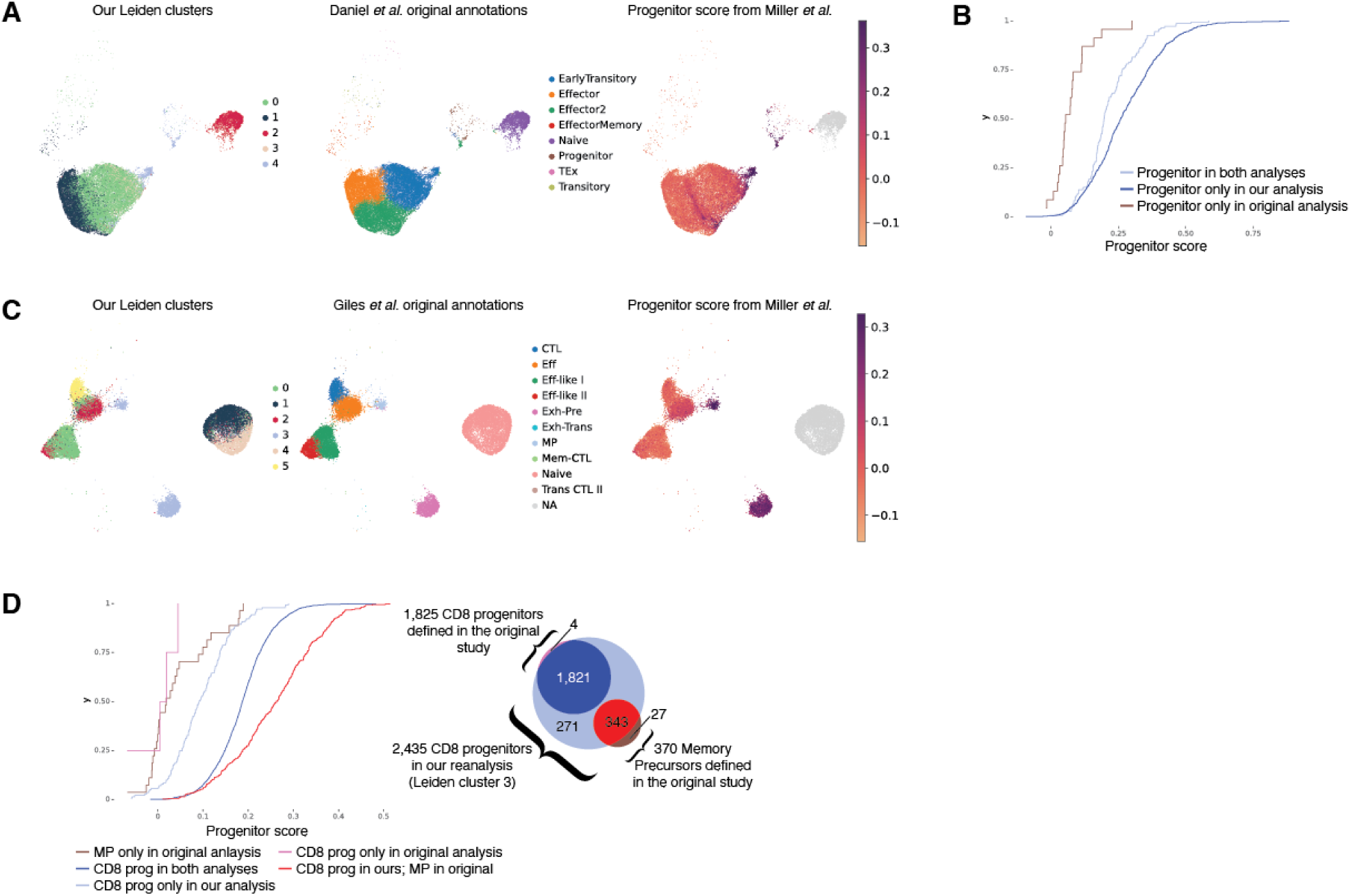
Additional analysis of the scATAC-seq data from Daniel *et al.* and Giles *et al.* (related to Figure 2). **(A-B)** Additional details of the reanalysis of the scATAC-seq data for CD8 T cells from Daniel *et al.*^35^ Complements the analysis in Fig. 2C-F. Leiden cluster 4 was annotated as progenitor CD8 T cells. **(A)** UMAP from the original paper, colored by: (*left* ) our Leiden clusters, (*center* ) the original annotations, and (*right* ) the epigenomic progenitor score from our bulk ATAC-seq compendium (Miller *et al.*). **(B)** CDF plot of progenitor scores for three groups: cells identified as progenitors in both analyses, cells in our Leiden cluster 4 not annotated as progenitors in the original paper, and cells annotated as progenitors in the original paper but not part of our cluster 4. The values shown in light blue or dark blue are significantly higher than brown (FDR < 3.9e*−*11, two-sided KS test). **(C-D)** Additional details of the reanalysis of the scATAC-seq data for CD8 T cells from Giles *et al.*^36^ Complements the analysis in Fig. 2G-J. Leiden cluster 3 was annotated as progenitor CD8 T cells, which included both memory precursors (MP) and progenitor cells (Pre-Exh) defined in the original publication. **(C)** UMAP from the original paper, colored by: (*left* ) our Leiden clusters, (*center* ) the original annotations, and (*right* ) the epigenomic progenitor score from our bulk ATAC-seq compendium. **(D)** CDF plot of the epigenomic progenitor scores for cells defined as progenitors in our annotations (Leiden cluster 3) or either progenitors or memory precursors (MPs) in the original paper. Similar to panel C. Colored lines for classifications are explained in the Venn diagram, including 271 cells shown in light blue that were identified as progenitors in our analysis but not identified as progenitors or MPs in the original publication (*right* ). The values shown in pink and brown are significantly lower than light blue (FDR < 4.5e*−*4, two-sided KS test).

**Fig. S8.**
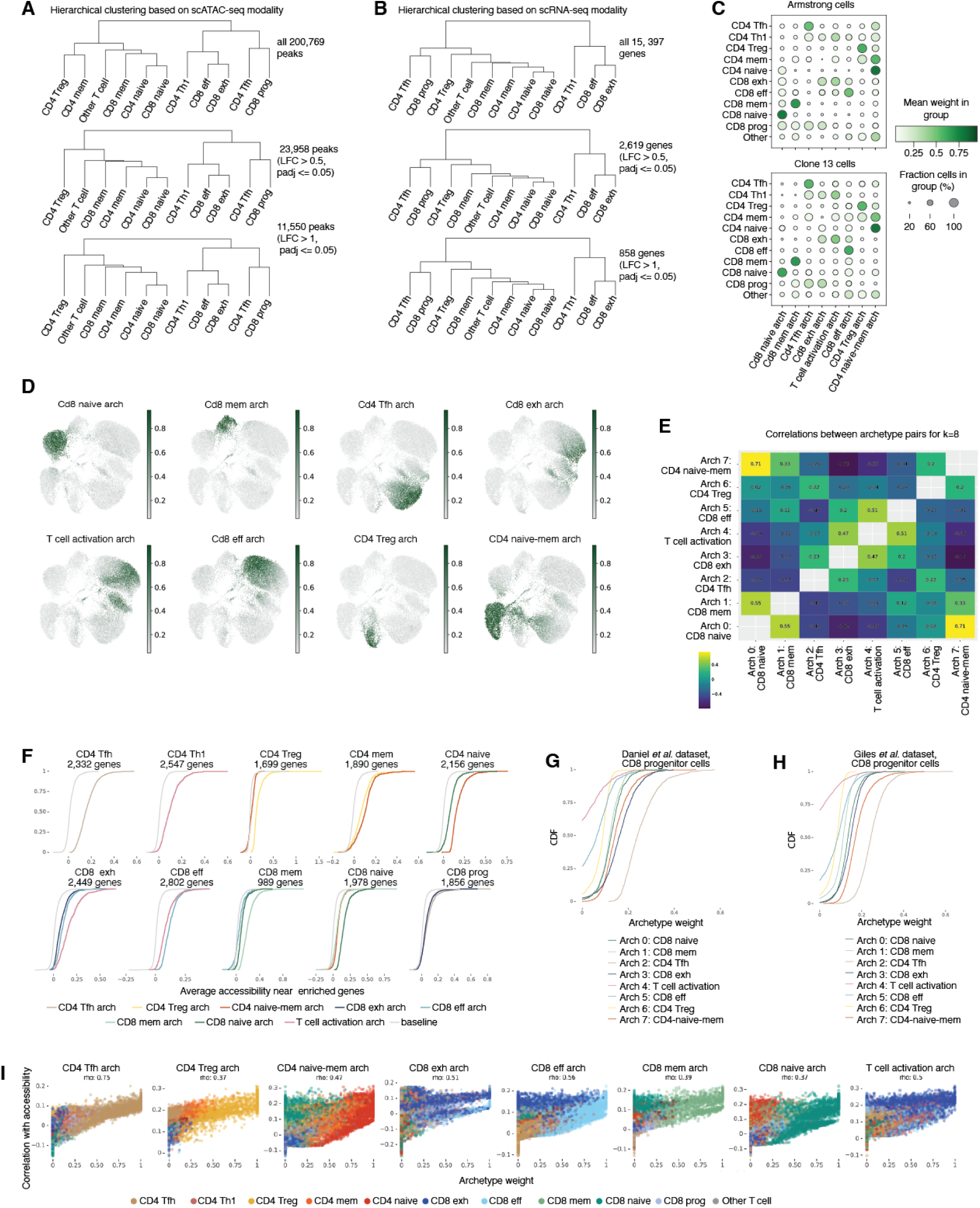
Archetypal analysis of the scATAC-seq data (related to Figure 3). **(A)** Hierarchical clustering of scATAC-seq data using different sets of peaks confirms similarity of chromatin accessibility between CD4 Tfh cells and progenitor CD8 T cells (Methods). **(B)** Hierarchical clustering of scRNA-seq data using different sets of genes confirms similarity of gene expression between CD4 Tfh cells and progenitor CD8 T cells. **(C)** Dotplots of archetypal weights for Armstrong and Clone 13 cells. Dot size represents the fraction of cells within a given cell state with strictly positive weight. The color of the dot indicates the average level of archetypal weight in that cell state. **(D)** UMAP plots showing cell-wise archetypal weights for each of the eight inferred archetypes. **(E)** Heatmap of pairwise correlations between archetypes, demonstrating that each archetype captures a distinct chromatin accessibility program. **(F)** CDF plots showing average accessibility of peaks near genes overexpressed in each cell type (LFC > 0, FDR *≤* 0.05) and associated with specific archetypes. Gray lines indicate baseline distributions of average accessibility for all peaks near a gene. Only the archetypes with significantly higher accessibility compared to baseline are shown for each cell state (KS test, FDR *≤* 0.05, and median accessibility difference *≥* 0.03). **(G)** CDF curves comparing the distribution of our archetype weights transferred to the Daniel *et al*. dataset, focusing on the CD8 progenitor population (Leiden cluster 4). Tfh archetype has the strongest weights. **(H)** CDF curves comparing the distribution of our archetype weights transferrred to the Giles *et al*. dataset, focusing on the CD8 progenitor population (Leiden cluster 3). Tfh archetype has the strongest weights. **(I)** Scatterplots showing the relationship between archetype weight and similarity of chromatin accessibility to each archetype. Each point represents one cell. The *x*-axis values represent the archetype weight in that cell. The *y*-axis values show the Spearman correlation between accessibility profile and the archetype peak scores over all peaks in that cell. Each cell is colored by its functional annotation, grouped by each of the eight archetypes.

**Fig. S9.**
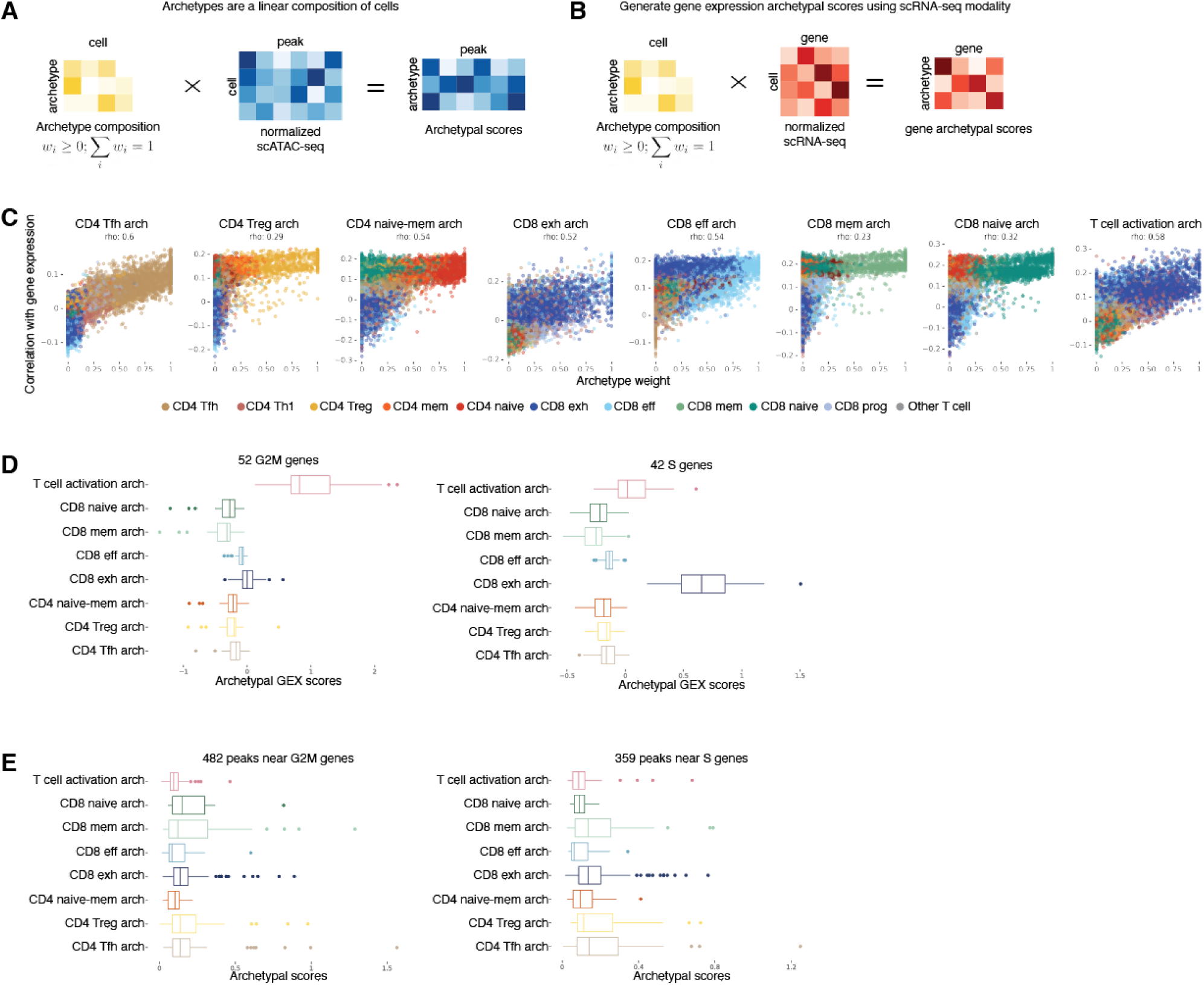
Gene expression patterns associated with scATAC-seq archetypes (related to Figure 3). **(A)** Schematic illustrating how the accessibility profile of each archetype (an archetypal score for each peak) is defined as a linear combination of cell accessibility profiles (see Methods). **(B)** Schematic showing how gene expression profiles for each archetype are derived using the cell weights from panel A. **(C)** Scatterplots showing the relationship between archetype weight and the similarity of gene expression to each archetype. Similar to Fig. S8I, but for gene expression instead of chromatin accessibility. Each point represents one cell. The *x*-axis values representing the archetype weight in that cell. The *y*-axis values show the Spearman correlation between the gene expression profile in that cell and the archetypal gene expression profile over all genes. Archetypal gene expression profiles were calculated using weighted average of cell expression profiles (see panel B, Methods). Each cell is colored by its functional annotation, grouped by each of the eight archetypes. **(D)** Boxplots showing gene archetypal scores for genes associated with cell cycle phases. Left: 52 genes related to the G2/M phase; right: 42 genes related to the S phase. **(E)** Boxplots showing chromatin accessibility archetypal scores for peaks near cell cycle-related genes. Left: 482 peaks near G2/M-phase-associated genes; right: 359 peaks near S-phase-associated genes.

**Fig. S10.**
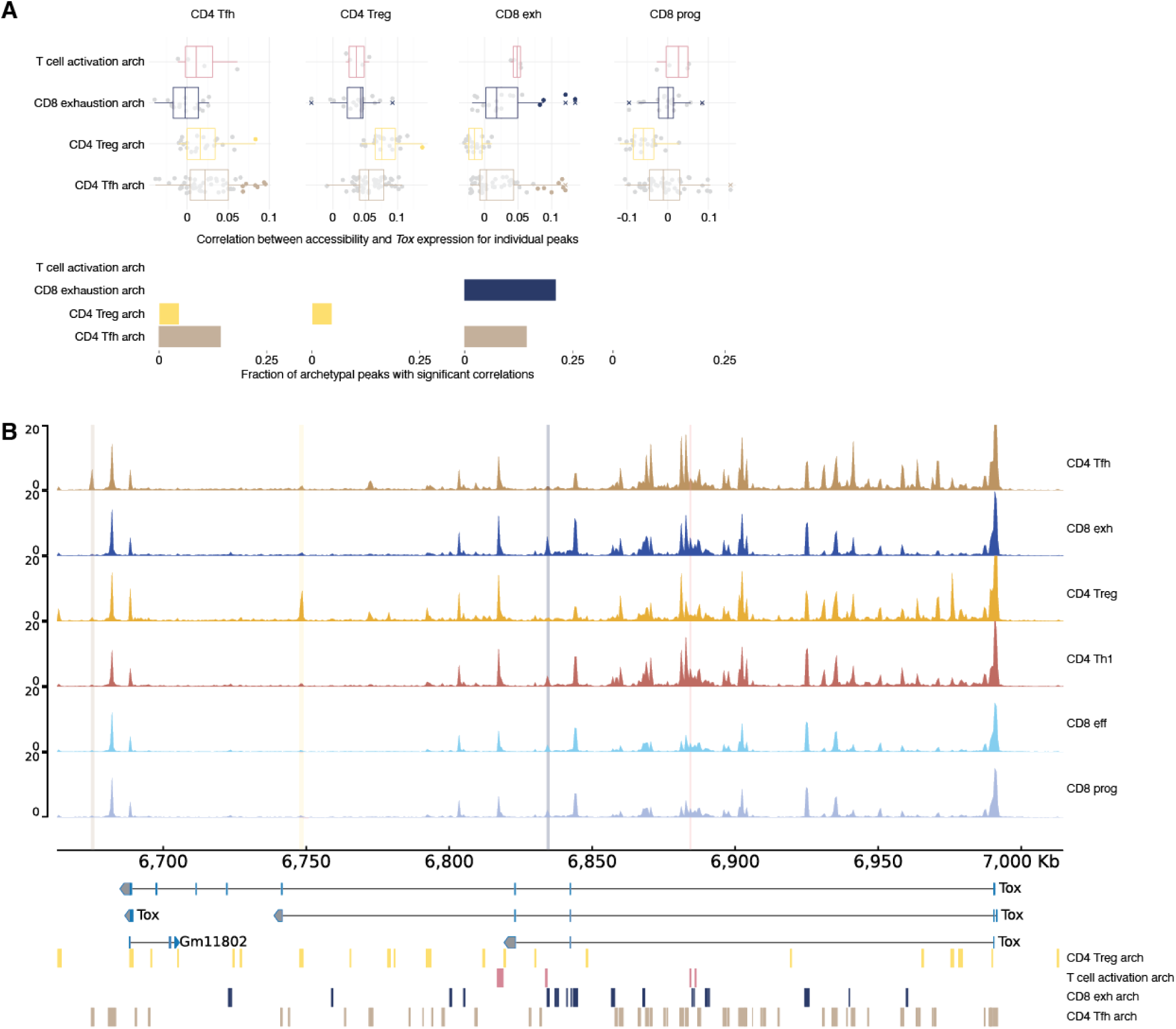
Archetypal analysis reveals cell state-specific regulation of *Tox* gene expression (related to Figure 3). **(A)** *Top*: Boxplot and scatterplot showing correlations between accessibility of individual peaks near *Tox* and *Tox* gene expression across different cell states. Peaks are grouped by their assigned scATAC-seq archetype. Significant correlations are colored by archetype; non-significant shown in gray. *Bottom*: Barplot showing the fraction of peaks within each archetype that significantly correlate with *Tox* expression in each cell population. **(B)** Pseudobulk scATAC-seq tracks around the *Tox* gene locus for different T cell subsets. Peaks are marked by rectangles at the bottom, colored by their archetype assignment. One representative peak with significant correlation is highlighted for each archetype.

**Fig. S11.**
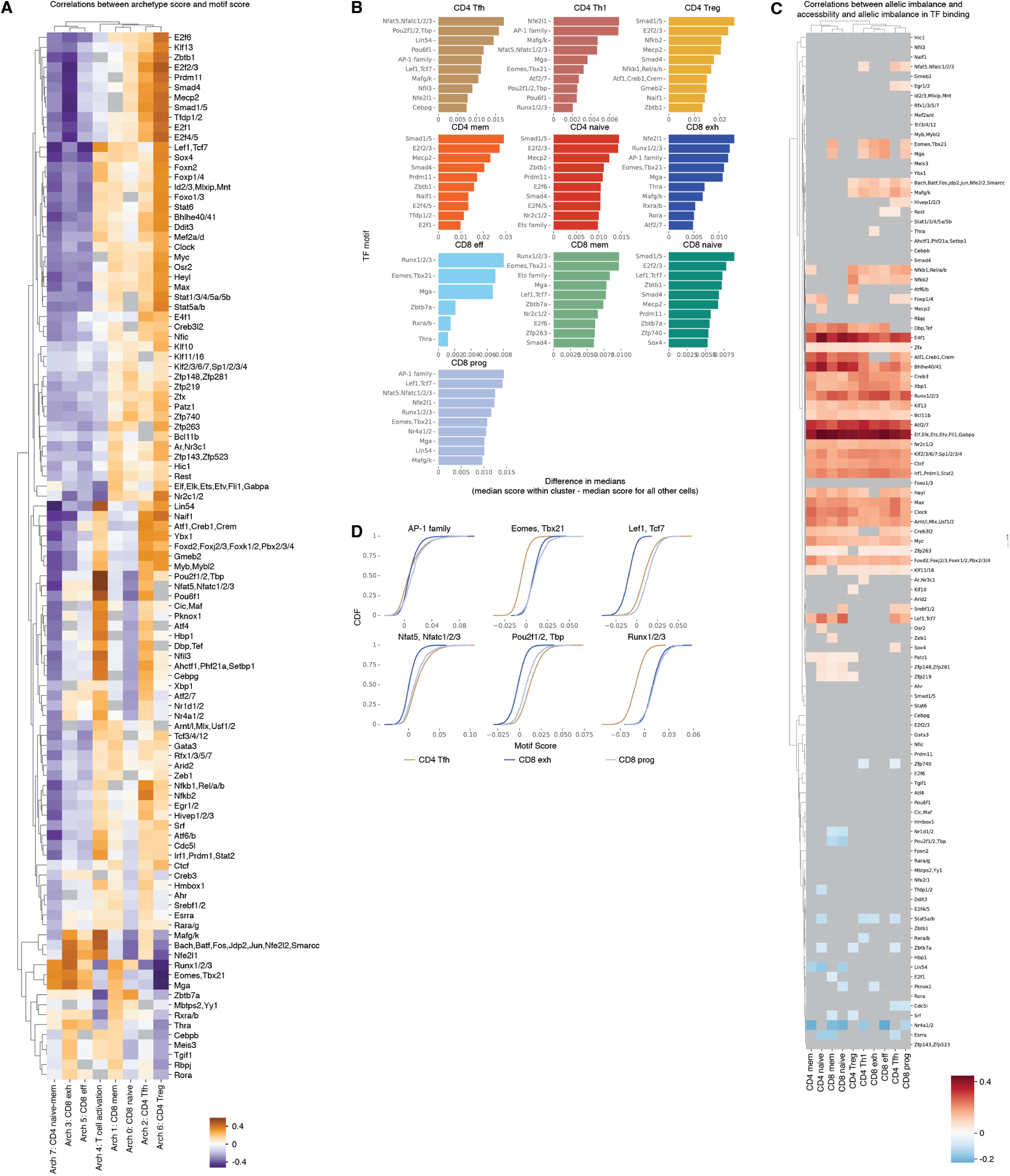
Transcription factors associated with distinct cell states and archetypes (related to Figure 4). **(A)** Clustered heatmap showing Spearman correlations between archetypal weights and motif scores. **(B)** Motifs most significantly enriched in each functional cell state compared with all other cells. The barplot shows the difference in medians (median motif score in cells of the selected cell state minus the median score for all other cells) for the top 10 motifs per cell state. All enrichments are significant (FDR *≤* 0.05, LFC > 0, 1-sided KS test, with Benjamini-Hochberg correction). **(C)** Clustered heatmap showing Spearman correlations between allelic imbalance of motif matches and chromatin accessibility across functional T cell states. Non-significant correlations (FDR > 0.05) are shown in gray. **(D)** CDF curves showing the motif score for selected motifs in three cell populations: CD4 Tfh cells, exhausted CD8 T cells, and progenitor CD8 T cells.

**Fig. S12.**
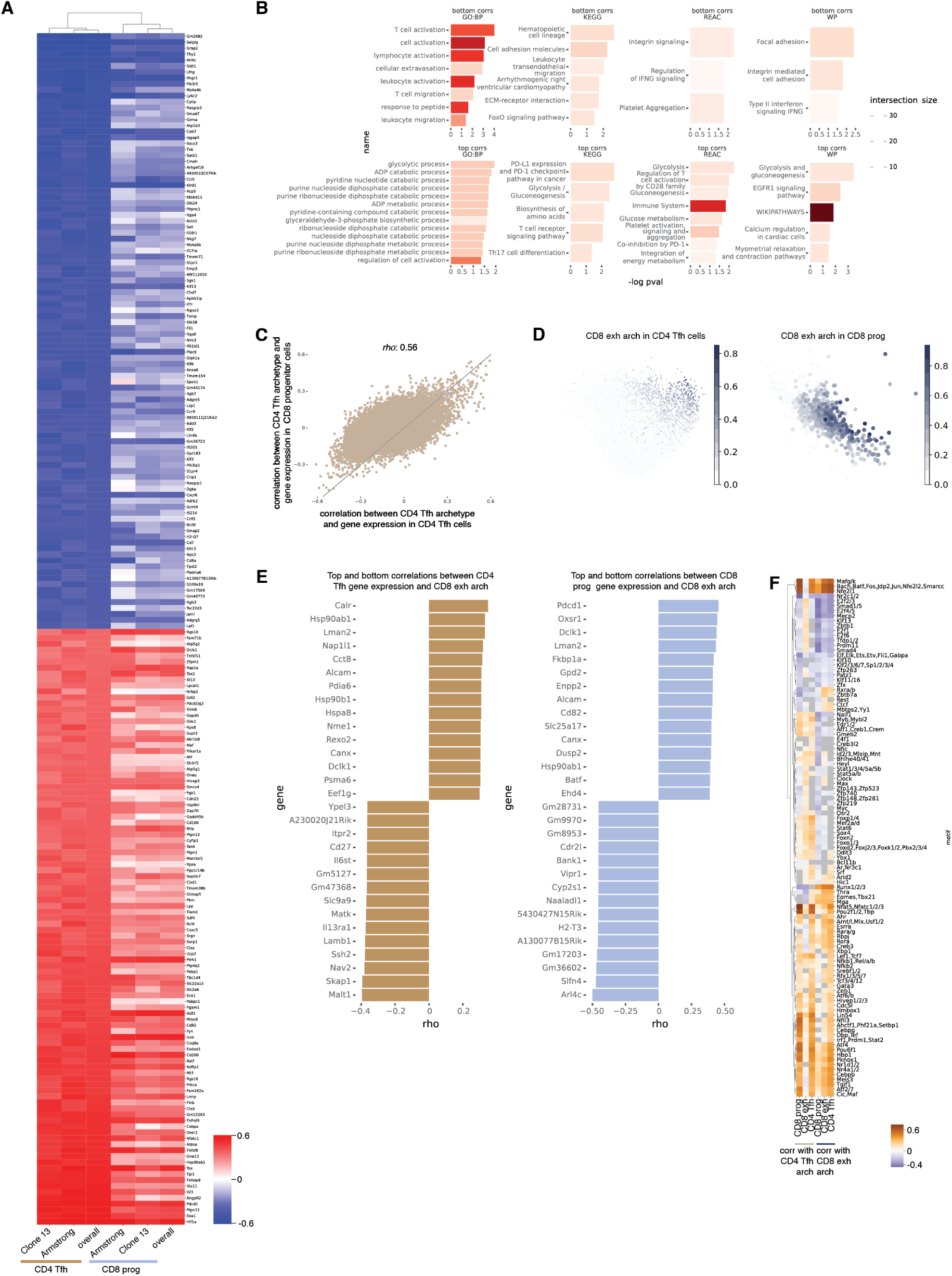
Tfh archetypal gradient in Tfh cells and progenitor Tcf1^+^ CD8 T cells (related to Figure 5). **(A)** Heatmap showing correlations between the Tfh archetype and gene expression in progenitor CD8 T cells and in CD4 Tfh cells. Genes shown are the top 100 most positively and top 100 most negatively correlated with Tfh archetypal weights, selected based on correlation across all Tfh cells. Correlations are computed separately for each cell state, as well as within Armstrong-only and Clone 13-only subsets. **(B)** Barplots showing significantly enriched pathways among the top 100 positively and top 100 negatively correlated genes with Tfh archetypal weights. Pathway databases include Gene Ontology (GO), KEGG, Reactome, and WikiPathways (see Methods). **(C)** Scatterplot comparing correlations between the CD4 Tfh archetype and gene expression levels in CD8 progenitor cells vs. CD4 Tfh cells. **(D)** UMAP highlighting the CD8 exhaustion archetypal weight in CD4 Tfh cells (*left* ) and progenitor CD8 T cells (*right* ). **(E)** Barplots showing the top 15 and bottom 15 genes significantly correlated with the CD8 exhaustion archetype, separately for CD4 Tfh cells and progenitor CD8 T cells. **(F)** Heatmap of correlations between archetype scores and transcription factor motif scores for all motifs, highlighting the CD4 Tfh and CD8 exhaustion archetypes across three selected cell types.

**Fig. S13.**
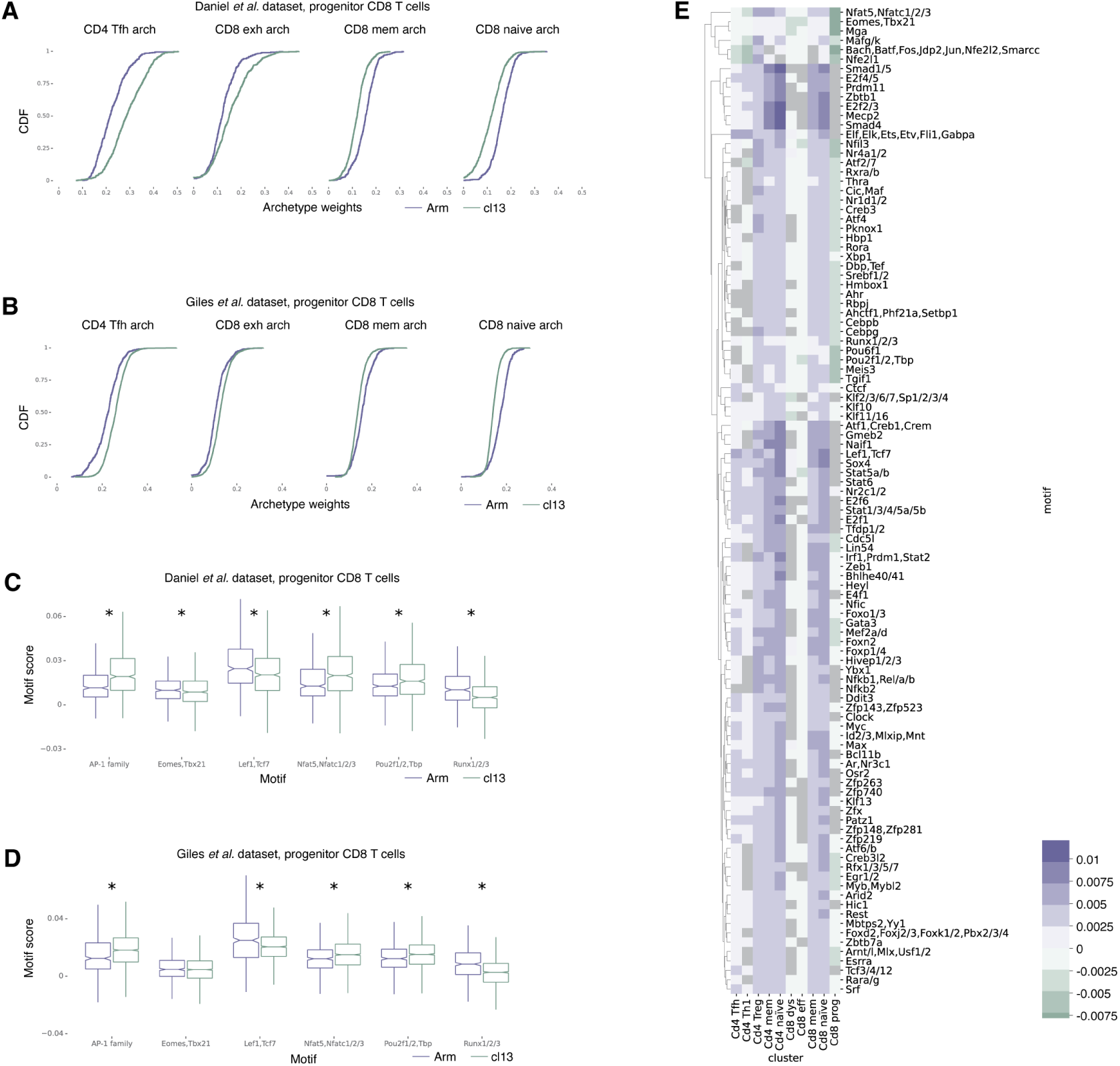
Exploring differences between Armstrong and Clone 13 cells (related to Figure 6). **(A)** CDF of our archetypal weights transferred to the Daniel *et al*. dataset, comparing between Armstrong and Clone 13 cells in the CD8 progenitor population (Leiden cluster 4). All distributions are significantly different (2-sided KS test, FDR < 2.4e*−*13) **(B)** CDF of our archetypal weights transferred to the Giles *et al*. dataset, comparing between Armstrong and Clone 13 cells in the CD8 progenitor population (Leiden cluster 3). All distributions are significantly different (2-sided KS test, FDR < 7.6e*−*19). **(C-D)** Boxplots comparing the motif scores of selected motifs between Armstrong and Clone 13 cells in the CD8 progenitor population in (C) the Daniel *et al*. dataset and (D) the Giles *et al*. dataset. Significant difference makred with asterisk (two-sided KS test, FDR *<* 0.05). **(E)** Heatmap comparing the motif scores of selected motifs between Armstrong and Clone 13 cells for all subpopulations in our dataset. Color represents the difference in medians between the Armstrong score and the Clone 13 score. Non-significant differences are shown in gray (FDR > 0.05, two-sided KS test).

## References

1. B. V. Kumar, T. J. Connors, and D. L. Farber. Human T cell development, localization, and function throughout life. Immunity, 48(2):202–213, 2018.

2. X. Fan and A. Y. Rudensky. Hallmarks of tissue-resident lymphocytes. Cell, 164(6):1198–1211, 2016.

3. H. Chi, M. Pepper, and P. G. Thomas. Principles and therapeutic applications of adaptive immunity. Cell, 187(9):2052–2078, 2024.

4. J. M. Schenkel and K. E. Pauken. Localization, tissue biology and T cell state—implications for cancer immunotherapy. Nature Reviews Immunology, 23(12):807–823, 2023.

5. M. Ruterbusch, K. B. Pruner, L. Shehata, and M. Pepper. In vivo CD4+ T cell differentiation and function: Revisiting the Th1/Th2 paradigm. Annual review of immunology, 38:705–725, 2020.

6. A. M. Miggelbrink, J. D. Jackson, S. J. Lorrey, E. S. Srinivasan, J. Waibl-Polania, D. S. Wilkinson, and P. E. Fecci. CD4 T-Cell Exhaustion: Does It Exist and What Are Its Roles in Cancer? Clinical Cancer Research: An Official Journal of the American Association for Cancer Research, 27(21):5742–5752, November 2021.

7. L. M. McLane, M. S. Abdel-Hakeem, and E. J. Wherry. CD8 T Cell Exhaustion During Chronic Viral Infection and Cancer. Annu Rev Immunol, 37:457–495, April 2019.

8. M. Hashimoto, A. O. Kamphorst, S. J. Im, H. T. Kissick, R. N. Pillai, S. S. Ramalingam, K. Araki, and R. Ahmed. CD8 T Cell Exhaustion in Chronic Infection and Cancer: Opportunities for Interventions. Annu Rev Med, 69(69):301–318, January 2018.

9. A. Kallies, D. Zehn, and D. T. Utzschneider. Precursor exhausted T cells: key to successful immunotherapy? Nature Reviews Immunology, 2019. PMID: 31591533.

10. S. Crotty. T Follicular Helper Cell Differentiation, Function, and Roles in Disease. Immunity, 41(4):529–542, October 2014.

11. Y. Kim, W. J. Greenleaf, and S. C. Bendall. Systems biology approaches to unravel lymphocyte subsets and function. Current opinion in immunology, 82:102323, 2023.

12. M. Efremova, R. Vento-Tormo, J.-E. Park, S. A. Teichmann, and K. R. James. Immunology in the era of single-cell technologies. Annual review of immunology, 38:727–757, 2020.

13. A. Giladi and I. Amit. Single-cell genomics: a stepping stone for future immunology discoveries. Cell, 172(1-2):14–21, 2018.

14. E. Kiner, E. Willie, B. Vijaykumar, K. Chowdhary, H. Schmutz, J. Chandler, A. Schnell, P. I. Thakore, G. LeGros, S. Mostafavi, et al. Gut CD4+ T cell phenotypes are a continuum molded by microbes, not by TH archetypes. Nature immunology, 22(2):216–228, 2021.

15. A. Wagner, A. Regev, and N. Yosef. Revealing the vectors of cellular identity with single-cell genomics. Nature biotechnology, 34(11):1145–1160, 2016.

16. S. Preissl, K. J. Gaulton, and B. Ren. Characterizing cis-regulatory elements using single-cell epigenomics. Nature Reviews Genetics, 24(1):21–43, 2023.

17. I. Eizenberg-Magar, J. Rimer, I. Zaretsky, D. Lara-Astiaso, S. Reich-Zeliger, and N. Friedman. Diverse continuum of CD4+ T-cell states is determined by hierarchical additive integration of cytokine signals. Proceedings of the National Academy of Sciences, 114(31):E6447–E6456, 2017.

18. M. R. Corces, J. D. Buenrostro, B. Wu, P. G. Greenside, S. M. Chan, J. L. Koenig, M. P. Snyder, J. K. Pritchard, A. Kundaje, W. J. Greenleaf, R. Majeti, and H. Y. Chang. Lineage-specific and single-cell chromatin accessibility charts human hematopoiesis and leukemia evolution. Nature Genetics, 48(10):1193–1203, October 2016.

19. H. K. Chung, B. McDonald, and S. M. Kaech. The architectural design of CD8+ T cell responses in acute and chronic infection: Parallel structures with divergent fates. Journal of Experimental Medicine, 218(4):e20201730, March 2021.

20. V. P. Badovinac, B. B. Porter, and J. T. Harty. Programmed contraction of CD8+ T cells after infection. Nature Immunology, 3(7):619–626, July 2002.

21. S. Walton, S. Mandaric, and A. Oxenius. CD4 T cell responses in latent and chronic viral infections. Frontiers in immunology, 4:105, 2013.

22. M. Andreatta, A. Tjitropranoto, Z. Sherman, M. C. Kelly, T. Ciucci, and S. J. Carmona. A CD4+ T cell reference map delineates subtype-specific adaptation during acute and chronic viral infections. Elife, 11:e76339, 2022.

23. T. Ciucci, M. S. Vacchio, Y. Gao, F. T. Ardori, J. Candia, M. Mehta, Y. Zhao, B. Tran, M. Pepper, L. Tessarollo, et al. The emergence and functional fitness of memory CD4+ T cells require the transcription factor Thpok. Immunity, 50(1):91–105, 2019.

24. R. Zander, M. Y. Kasmani, Y. Chen, P. Topchyan, J. Shen, S. Zheng, R. Burns, J. Ingram, C. Cui, N. Joshi, et al. Tfh-cell-derived interleukin 21 sustains effector CD8+ T cell responses during chronic viral infection. Immunity, 55(3):475–493, 2022.

25. M. Sade-Feldman, K. Yizhak, S. L. Bjorgaard, J. P. Ray, C. G. de Boer, R. W. Jenkins, D. J. Lieb, J. H. Chen, D. T. Frederick, M. Barzily-Rokni, et al. Defining T cell states associated with response to checkpoint immunotherapy in melanoma. Cell, 175(4):998–1013, 2018.

26. C. U. Blank, W. N. Haining, W. Held, P. G. Hogan, A. Kallies, E. Lugli, R. C. Lynn, M. Philip, A. Rao, N. P. Restifo, et al. Defining ‘T cell exhaustion’. Nature Reviews Immunology, 19(11):665–674, 2019.

27. B. C. Miller, D. R. Sen, R. Al Abosy, K. Bi, Y. V. Virkud, M. W. LaFleur, K. B. Yates, A. Lako, K. Felt, G. S. Naik, M. Manos, E. Gjini, J. R. Kuchroo, J. J. Ishizuka, J. L. Collier, G. K. Griffin, S. Maleri, D. E. Comstock, S. A. Weiss, F. D. Brown, A. Panda, M. D. Zimmer, R. T. Manguso, F. S. Hodi, S. J. Rodig, A. H. Sharpe, and W. N. Haining. Subsets of exhausted CD8+ T cells differentially mediate tumor control and respond to checkpoint blockade. Nature Immunology, 20(3):326–336, March 2019.

28. I. Siddiqui, K. Schaeuble, V. Chennupati, S. A. Fuertes Marraco, S. Calderon-Copete, D. Pais Ferreira, S. J. Carmona, L. Scarpellino, D. Gfeller, S. Pradervand, S. A. Luther, D. E. Speiser, and W. Held. Intratumoral Tcf1+PD-1+CD8+ T Cells with Stem-like Properties Promote Tumor Control in Response to Vaccination and Checkpoint Blockade Immunotherapy. Immunity, 50(1):195–211.e10, January 2019.

29. A. Kallies, D. Zehn, and D. T. Utzschneider. Precursor exhausted T cells: key to successful immunotherapy? Nature Reviews Immunology, 20(2):128–136, February 2020.

30. Z. Fang, X. Ding, H. Huang, H. Jiang, J. Jiang, and X. Zheng. Revolutionizing tumor immunotherapy: unleashing the power of progenitor exhausted T cells. Cancer Biology Medicine, May 2024.

31. J. R. Giles, A.-M. Globig, S. M. Kaech, and E. J. Wherry. CD8+ T cells in the cancer-immunity cycle. Immunity, 56(10):2231–2253, 2023.

32. Y. Pritykin, J. van der Veeken, A. R. Pine, Y. Zhong, M. Sahin, L. Mazutis, D. Pe’er, A. Y. Rudensky, and C. S. Leslie. A unified atlas of CD8 T cell dysfunctional states in cancer and infection. Molecular Cell, 81(11):2477–2493.e10, 2021.

33. N. S. Joshi, W. Cui, A. Chandele, H. K. Lee, D. R. Urso, J. Hagman, L. Gapin, and S. M. Kaech. Inflammation Directs Memory Precursor and Short-Lived Effector CD8+ T Cell Fates via the Graded Expression of T-bet Transcription Factor. Immunity, 27(2):281–295, August 2007.

34. S. M. Kaech and W. Cui. Transcriptional control of effector and memory CD8+ T cell differentiation. Nature Reviews Immunology, 12(11):749–761, November 2012.

35. B. Daniel, K. E. Yost, S. Hsiung, K. Sandor, Y. Xia, Y. Qi, K. J. Hiam-Galvez, M. Black, C. J. Raposo, Q. Shi, S. L. Meier, J. A. Belk, J. R. Giles, E. J. Wherry, H. Y. Chang, T. Egawa, and A. T. Satpathy. Divergent clonal differentiation trajectories of T cell exhaustion. Nature Immunology, 23(11):1614–1627, November 2022.

36. J. R. Giles, S. F. Ngiow, S. Manne, A. E. Baxter, O. Khan, P. Wang, R. Staupe, M. S. Abdel-Hakeem, H. Huang, D. Mathew, M. M. Painter, J. E. Wu, Y. J. Huang, R. R. Goel, P. K. Yan, G. C. Karakousis, X. Xu, T. C. Mitchell, A. C. Huang, and E. J. Wherry. Shared and distinct biological circuits in effector, memory and exhausted CD8+ T cells revealed by temporal single-cell transcriptomics and epigenetics. Nature Immunology, 23(11):1600–1613, November 2022.

37. D. T. Utzschneider, S. S. Gabriel, D. Chisanga, R. Gloury, P. M. Gubser, A. Vasanthakumar, W. Shi, and A. Kallies. Early precursor T cells establish and propagate T cell exhaustion in chronic infection. Nature Immunology, 21(10):1256–1266, October 2020.

38. T. Chu, M. Wu, B. Hoellbacher, G. P. de Almeida, C. Wurmser, J. Berner, L. V. Donhauser, A.-K. Gerullis, S. Lin, J. D. Cepeda-Mayorga, I. I. Kilb, L. Bongers, F. Toppeta, P. Strobl, B. Youngblood, A. M. Schulz, A. Zippelius, P. A. Knolle, M. Heinig, C.-P. Hackstein, and D. Zehn. Precursors of exhausted T cells are pre-emptively formed in acute infection. Nature, 640(8059):782–792, April 2025.

39. A. D. T. McManus, R. M. Valanparambil, C. B. Medina, C. D. Scharer, D. J. McGuire, E. Sobierajska, Y. Hu, D. Y. Chang, A. Wieland, J. Lee, T. H. Nasti, M. Hashimoto, J. L. Ross, N. Prokhnevska, M. A. Cardenas, A. L. Gill, E. C. Clark, K. Abadie, A. J. Kumar, J. Kaye, B. B. Au-Yeung, H. Y. Kueh, H. T. Kissick, and R. Ahmed. An early precursor CD8+ T cell that adapts to acute or chronic viral infection. Nature, 640(8059):772–781, April 2025.

39. C. Gago da Graça, A. A. Sheikh, D. M. Newman, L. Wen, S. Li, J. Shen, Y. Zhang, S. S. Gabriel, D. Chisanga, J. Seow, A. Poch, L. Rausch, M.-H. T. Nguyen, J. Singh, C.-H. Su, L. A. Cluse, C. Tsui, T. N. Burn, S. L. Park, B. Von Scheidt, L. K. Mackay, A. Vasanthakumar, D. Bending, W. Shi, W. Cui, J. Schröder, R. W. Johnstone, A. Kallies, and D. T. Utzschneider. Stem-like memory and precursors of exhausted T cells share a common progenitor defined by ID3 expression. Science Immunology, 10(103):eadn1945, January 2025.

41. A. S. J. Im, M. Hashimoto, M. Y. Gerner, J. Lee, H. T. Kissick, M. C. Burger, Q. Shan, J. S. Hale, J. Lee, T. H. Nasti, A. H. Sharpe, G. J. Freeman, R. N. Germain, H. I. Nakaya, H.-H. Xue, and R. Ahmed. Defining CD8+ T cells that provide the proliferative burst after PD-1 therapy. Nature, 537(7620):417–421, September 2016.

40. W. D. Green, A. Gomez, A. L. Plotkin, B. M. Pratt, E. F. Merritt, G. N. Mullins, N. P. Kren, J. L. Modliszewski, V. Zhabotynsky, M. G. Woodcock, et al. Enhancer-driven gene regulatory networks reveal transcription factors governing T cell adaptation and differentiation in the tumor microenvironment. Immunity, 2025.

41. D. Ucar, E. J. Márquez, C.-H. Chung, R. Marches, R. J. Rossi, A. Uyar, T.-C. Wu, J. George, M. L. Stitzel, A. K. Palucka, et al. The chromatin accessibility signature of human immune aging stems from CD8+ T cells. Journal of Experimental Medicine, 214(10):3123–3144, 2017.

42. Y. Zhong, S. K. Walker, Y. Pritykin, C. S. Leslie, A. Y. Rudensky, and J. van der Veeken. Hierarchical regulation of the resting and activated T cell epigenome by major transcription factor families. Nature immunology, 23:122–134, 2022.

43. G. P. Mognol, R. Spreafico, V. Wong, J. P. Scott-Browne, S. Togher, A. Hoffmann, P. G. Hogan, A. Rao, and S. Trifari. Exhaustion-associated regulatory regions in CD8+ tumor-infiltrating T cells. Proceedings of the National Academy of Sciences, 114(13):E2776–E2785, March 2017.

44. C. D. Scharer, A. P. R. Bally, B. Gandham, and J. M. Boss. Cutting Edge: Chromatin Accessibility Programs CD8 T Cell Memory. The Journal of Immunology, 198(6):2238–2243, March 2017.

45. D. R. Sen, J. Kaminski, R. A. Barnitz, M. Kurachi, U. Gerdemann, K. B. Yates, H.-W. Tsao, J. Godec, M. W. LaFleur, F. D. Brown, et al. The epigenetic landscape of T cell exhaustion. Science, 354(6316):1165–1169, 2016.

46. S. Ma, B. Zhang, L. M. LaFave, A. S. Earl, Z. Chiang, Y. Hu, J. Ding, A. Brack, V. K. Kartha, T. Tay, et al. Chromatin potential identified by shared single-cell profiling of RNA and chromatin. Cell, 183(4):1103–1116, 2020.

47. A. J. González, M. Setty, and C. S. Leslie. Early enhancer establishment and regulatory locus complexity shape transcriptional programs in hematopoietic differentiation. Nature genetics, 47(11):1249–1259, 2015.

48. E. Fagerberg, J. Attanasio, C. Dien, J. Singh, E. A. Kessler, L. Abdullah, J. Shen, B. G. Hunt, K. A. Connolly, E. De Brouwer, et al. KLF2 maintains lineage fidelity and suppresses CD8 T cell exhaustion during acute LCMV infection. Science, 387(6735):eadn2337, 2025.

49. J. Chen, I. F. López-Moyado, H. Seo, C.-W. J. Lio, L. J. Hempleman, T. Sekiya, A. Yoshimura, J. P. Scott-Browne, and A. Rao. NR4A transcription factors limit CAR T cell function in solid tumours. Nature, 567(7749):530–534, March 2019.

50. B. C. Miller, D. R. Sen, R. Al Abosy, K. Bi, Y. V. Virkud, M. W. LaFleur, K. B. Yates, A. Lako, K. Felt, G. S. Naik, M. Manos, E. Gjini, J. R. Kuchroo, J. J. Ishizuka, J. L. Collier, G. K. Griffin, S. Maleri, D. E. Comstock, S. A. Weiss, F. D. Brown, A. Panda, M. D. Zimmer, R. T. Manguso, F. S. Hodi, S. J. Rodig, A. H. Sharpe, and W. N. Haining. Subsets of exhausted CD8+ T cells differentially mediate tumor control and respond to checkpoint blockade. Nature Immunology, 20(3):326–336, March 2019.

51. A. Cutler, , and L. Breiman. Archetypal Analysis. Technometrics, 36(4):338–347, November 1994.

52. M. Mørup and L. K. Hansen. Archetypal analysis for machine learning and data mining. Neurocomputing, 80:54–63, March 2012.

53. A. Damle, , and Y. Sun. A Geometric Approach to Archetypal Analysis and Nonnegative Matrix Factorization. Technometrics, 59(3):361–370, July 2017.

54. H. Niu and H. Wang. TOX regulates T lymphocytes differentiation and its function in tumor. Frontiers in Immunology, 14, March 2023.

55. O. Khan, J. R. Giles, S. McDonald, S. Manne, S. F. Ngiow, K. P. Patel, M. T. Werner, A. C. Huang, K. A. Alexander, J. E. Wu, J. Attanasio, P. Yan, S. M. George, B. Bengsch, R. P. Staupe, G. Donahue, W. Xu, R. K. Amaravadi, X. Xu, G. C. Karakousis, T. C. Mitchell, L. M. Schuchter, J. Kaye, S. L. Berger, and E. J. Wherry. TOX transcriptionally and epigenetically programs CD8+ T cell exhaustion. Nature, 571(7764):211–218, July 2019.

56. F. Alfei, K. Kanev, M. Hofmann, M. Wu, H. E. Ghoneim, P. Roelli, D. T. Utzschneider, M. von Hoesslin, J. G. Cullen, Y. Fan, V. Eisenberg, D. Wohlleber, K. Steiger, D. Merkler, M. Delorenzi, P. A. Knolle, C. J. Cohen, R. Thimme, B. Youngblood, and D. Zehn. TOX reinforces the phenotype and longevity of exhausted T cells in chronic viral infection. Nature, 571(7764):265–269, July 2019.

57. C. Yao, H.-W. Sun, N. E. Lacey, Y. Ji, E. A. Moseman, H.-Y. Shih, E. F. Heuston, M. Kirby, S. Anderson, J. Cheng, O. Khan, R. Handon, J. Reilley, J. Fioravanti, J. Hu, S. Gossa, E. J. Wherry, L. Gattinoni, D. B. McGavern, J. J. O’Shea, P. L. Schwartzberg, and T. Wu. Single-cell RNA-seq reveals TOX as a key regulator of CD8+ T cell persistence in chronic infection. Nature Immunology, 20(7):890–901, July 2019.

58. A. C. Scott, F. Dündar, P. Zumbo, S. S. Chandran, C. A. Klebanoff, M. Shakiba, P. Trivedi, L. Menocal, H. Appleby, S. Camara, D. Zamarin, T. Walther, A. Snyder, M. R. Femia, E. A. Comen, H. Y. Wen, M. D. Hellmann, N. Anandasabapathy, Y. Liu, N. K. Altorki, P. Lauer, O. Levy, M. S. Glickman, J. Kaye, D. Betel, M. Philip, and A. Schietinger. TOX is a critical regulator of tumour-specific T cell differentiation. Nature, 571(7764):270–274, July 2019.

59. J. van der Veeken, A. Glasner, Y. Zhong, W. Hu, Z.-M. Wang, R. Bou-Puerto, L.-M. Charbonnier, T. A. Chatila, C. S. Leslie, and A. Y. Rudensky. The transcription factor Foxp3 shapes regulatory T cell identity by tuning the activity of trans-acting intermediaries. Immunity, 53(5):971–984, 2020.

60. G. J. Martinez, J. K. Hu, R. M. Pereira, J. S. Crampton, S. Togher, N. Bild, S. Crotty, and A. Rao. Cutting Edge: NFAT Transcription Factors Promote the Generation of Follicular Helper T Cells in Response to Acute Viral Infection. The Journal of Immunology, 196(5):2015–2019, March 2016.

61. G. J. Martinez, R. M. Pereira, T. Äijö, E. Y. Kim, F. Marangoni, M. E. Pipkin, S. Togher, V. Heissmeyer, Y. C. Zhang, S. Crotty, E. D. Lamperti, K. M. Ansel, T. R. Mempel, H. Lähdesmäki, P. G. Hogan, and A. Rao. The Transcription Factor NFAT Promotes Exhaustion of Activated CD8+ T Cells. Immunity, 42(2):265–278, February 2015.

62. X. Zhang, C. Zhang, M. Qiao, C. Cheng, N. Tang, S. Lu, W. Sun, B. Xu, Y. Cao, X. Wei, Y. Wang, W. Han, and H. Wang. Depletion of BATF in CAR-T cells enhances antitumor activity by inducing resistance against exhaustion and formation of central memory cells. Cancer Cell, 40(11):1407–1422.e7, November 2022.

63. H.-W. Tsao, J. Kaminski, M. Kurachi, R. A. Barnitz, M. A. DiIorio, M. W. LaFleur, W. Ise, T. Kurosaki, E. J. Wherry, W. N. Haining, et al. Batf-mediated epigenetic control of effector CD8+ T cell differentiation. Science immunology, 7(68):eabi4919, 2022.

64. H. Seo, J. Chen, E. González-Avalos, D. Samaniego-Castruita, A. Das, Y. H. Wang, I. F. López-Moyado, R. O. Georges, W. Zhang, A. Onodera, et al. TOX and TOX2 transcription factors cooperate with NR4A transcription factors to impose CD8+ T cell exhaustion. Proceedings of the National Academy of Sciences, 116(25):12410–12415, 2019.

65. J.-C. Beltra, S. Manne, M. S. Abdel-Hakeem, M. Kurachi, J. R. Giles, Z. Chen, V. Casella, S. F. Ngiow, O. Khan, Y. J. Huang, P. Yan, K. Nzingha, W. Xu, R. K. Amaravadi, X. Xu, G. C. Karakousis, T. C. Mitchell, L. M. Schuchter, A. C. Huang, and E. J. Wherry. Developmental Relationships of Four Exhausted CD8+ T Cell Subsets Reveals Underlying Transcriptional and Epigenetic Landscape Control Mechanisms. Immunity, 52(5):825–841.e8, May 2020.

66. D. Zehn, R. Thimme, E. Lugli, G. P. de Almeida, and A. Oxenius. ‘Stem-like’precursors are the fount to sustain persistent CD8+ T cell responses. Nature immunology, 23(6):836–847, 2022.

67. Y. Lv, L. Ricard, B. Gaugler, H. Huang, and Y. Ye. Biology and clinical relevance of follicular cytotoxic T cells. Frontiers in Immunology, 13:1036616, 2022.

68. R. He, S. Hou, C. Liu, A. Zhang, Q. Bai, M. Han, Y. Yang, G. Wei, T. Shen, X. Yang, L. Xu, X. Chen, Y. Hao, P. Wang, C. Zhu, J. Ou, H. Liang, T. Ni, X. Zhang, X. Zhou, K. Deng, Y. Chen, Y. Luo, J. Xu, H. Qi, Y. Wu, and L. Ye. Follicular CXCR5-expressing CD8+ T cells curtail chronic viral infection. Nature, 537(7620):412–416, September 2016.

69. Y. A. Leong, Y. Chen, H. S. Ong, D. Wu, K. Man, C. Deleage, M. Minnich, B. J. Meckiff, Y. Wei, Z. Hou, D. Zotos, K. A. Fenix, A. Atnerkar, S. Preston, J. G. Chipman, G. J. Beilman, C. C. Allison, L. Sun, P. Wang, J. Xu, J. G. Toe, H. K. Lu, Y. Tao, U. Palendira, A. L. Dent, A. L. Landay, M. Pellegrini, I. Comerford, S. R. McColl, T. W. Schacker, H. M. Long, J. D. Estes, M. Busslinger, G. T. Belz, S. R. Lewin, A. Kallies, and D. Yu. CXCR5+ follicular cytotoxic T cells control viral infection in B cell follicles. Nature Immunology, 17(10):1187–1196, October 2016.

70. Y. Chen, M. Yu, Y. Zheng, G. Fu, G. Xin, W. Zhu, L. Luo, R. Burns, Q.-Z. Li, A. L. Dent, et al. CXCR5+ PD-1+ follicular helper CD8 T cells control B cell tolerance. Nature communications, 10(1):4415, 2019.

71. D. Yu and L. Ye. A portrait of CXCR5+ follicular cytotoxic CD8+ T cells. Trends in immunology, 39(12):965–979, 2018.

72. Z. Chen, Z. Ji, S. F. Ngiow, S. Manne, Z. Cai, A. C. Huang, J. Johnson, R. P. Staupe, B. Bengsch, C. Xu, S. Yu, M. Kurachi, R. S. Herati, L. A. Vella, A. E. Baxter, J. E. Wu, O. Khan, J.-C. Beltra, J. R. Giles, E. Stelekati, L. M. McLane, C. W. Lau, X. Yang, S. L. Berger, G. Vahedi, H. Ji, and E. J. Wherry. TCF-1-Centered Transcriptional Network Drives an Effector versus Exhausted CD8 T Cell-Fate Decision. Immunity, 51(5):840–855.e5, November 2019.

73. K. Kanev, M. Wu, A. Drews, P. Roelli, C. Wurmser, M. von Hösslin, and D. Zehn. Proliferation-competent Tcf1+ CD8 T cells in dysfunctional populations are CD4 T cell help independent. Proceedings of the National Academy of Sciences, 116(40):20070–20076, 2019.

74. V. van der Heide, G. Laghlali, B. Davenport, B. Cubitt, V. Roudko, D. Choo, K. Jhun, E. Humblin, A. Vaidya, K. Angeliadis, et al. Prolonged but finite antigen presentation promotes reversible defects of “helpless” memory CD8+ T cells. Immunity, 2025.

75. C. Manes, M. G. Moreno, J. Cimons, M. A. D’Antonio, T. M. Brunetti, M. G. Harbell, S. Selva, C. Mizenko, T. L. Borko, E. L. Lasda, et al. B cells shape naive CD8+ T cell programming. The Journal of Clinical Investigation, 135(12), 2025.

76. M. L. Burger, A. M. Cruz, G. E. Crossland, G. Gaglia, C. C. Ritch, S. E. Blatt, A. Bhutkar, D. Canner, T. Kienka, S. Z. Tavana, A. L. Barandiaran, A. Garmilla, J. M. Schenkel, M. Hillman, I. de los Rios Kobara, A. Li, A. M. Jaeger, W. L. Hwang, P. M. K. Westcott, M. P. Manos, M. M. Holovatska, F. S. Hodi, A. Regev, S. Santagata, and T. Jacks. Antigen dominance hierarchies shape TCF1+ progenitor CD8 T cell phenotypes in tumors. Cell, 184(19):4996–5014.e26, September 2021.

77. J. M. Schenkel, R. H. Herbst, D. Canner, A. Li, M. Hillman, S.-L. Shanahan, G. Gibbons, O. C. Smith, J. Y. Kim, P. Westcott, W. L. Hwang, W. A. Freed-Pastor, G. Eng, M. S. Cuoco, P. Rogers, J. K. Park, M. L. Burger, O. Rozenblatt-Rosen, L. Cong, K. E. Pauken, A. Regev, and T. Jacks. Conventional type I dendritic cells maintain a reservoir of proliferative tumor-antigen specific TCF-1+ CD8+ T cells in tumor-draining lymph nodes. Immunity, 54(10):2338–2353.e6, October 2021.

78. S. Nakandakari-Higa, S. Walker, M. C. C. Canesso, V. van der Heide, A. Chudnovskiy, D.-Y. Kim, J. T. Jacobsen, R. Parsa, J. Bilanovic, S. M. Parigi, K. Fiedorczuk, E. Fuchs, A. M. Bilate, G. Pasqual, D. Mucida, A. O. Kamphorst, Y. Pritykin, and G. D. Victora. Universal recording of immune cell interactions in vivo. Nature, 627:399–406, 2024.

79. B. Langmead and S. L. Salzberg. Fast gapped-read alignment with Bowtie 2. Nature Methods, 9(4):357–359, April 2012.

80. P. Danecek, J. K. Bonfield, J. Liddle, J. Marshall, V. Ohan, M. O. Pollard, A. Whitwham, T. Keane, S. A. McCarthy, R. M. Davies, and H. Li. Twelve years of SAMtools and BCFtools. GigaScience, 10(2):giab008, February 2021.

81. A. R. Quinlan and I. M. Hall. BEDTools: a flexible suite of utilities for comparing genomic features. Bioinformatics, 26(6):841–842, March 2010.

82. Y. Zhang, T. Liu, C. A. Meyer, J. Eeckhoute, D. S. Johnson, B. E. Bernstein, C. Nusbaum, R. M. Myers, M. Brown, W. Li, and X. S. Liu. Model-based Analysis of ChIP-Seq (MACS). Genome Biology, 9(9):R137, September 2008.

83. Q. Li, J. B. Brown, H. Huang, and P. J. Bickel. Measuring reproducibility of high-throughput experiments. The Annals of Applied Statistics, 5(3):1752–1779, September 2011.

84. J. M. Granja, M. R. Corces, S. E. Pierce, S. T. Bagdatli, H. Choudhry, H. Y. Chang, and W. J. Greenleaf. ArchR is a scalable software package for integrative single-cell chromatin accessibility analysis. Nature Genetics, 53(3):403–411, March 2021.

85. I. Korsunsky, N. Millard, J. Fan, K. Slowikowski, F. Zhang, K. Wei, Y. Baglaenko, M. Brenner, P.-r. Loh, and S. Raychaudhuri. Fast, sensitive and accurate integration of single-cell data with Harmony. Nature Methods, 16(12):1289–1296, December 2019.

86. M. Lawrence, W. Huber, H. Pagès, P. Aboyoun, M. Carlson, R. Gentleman, M. T. Morgan, and V. J. Carey. Software for Computing and Annotating Genomic Ranges. PLOS Computational Biology, 9(8):e1003118, August 2013.

87. M. I. Love, W. Huber, and S. Anders. Moderated estimation of fold change and dispersion for RNA-seq data with DESeq2. Genome Biology, 15(12):550, December 2014.

88. F. A. Wolf, P. Angerer, and F. J. Theis. SCANPY: large-scale single-cell gene expression data analysis. Genome Biology, 19(1):15, February 2018.

89. L. D. Martens, D. S. Fischer, V. A. Yépez, F. J. Theis, and J. Gagneur. Modeling fragment counts improves single-cell ATAC-seq analysis. Nature Methods, 21(1):28–31, January 2024.

90. J. Lause, P. Berens, and D. Kobak. Analytic Pearson residuals for normalization of single-cell RNA-seq UMI data. Genome Biology, 22(1):258, September 2021.

91. S. L. Wolock, R. Lopez, and A. M. Klein. Scrublet: Computational Identification of Cell Doublets in Single-Cell Transcriptomic Data. Cell Systems, 8(4):281–291.e9, April 2019.

92. W. J. Kent, A. S. Zweig, G. Barber, A. S. Hinrichs, and D. Karolchik. BigWig and BigBed: enabling browsing of large distributed datasets. Bioinformatics, 26(17):2204–2207, September 2010.

93. S. Lianoglou, V. Garg, J. L. Yang, C. S. Leslie, and C. Mayr. Ubiquitously transcribed genes use alternative polyadenylation to achieve tissue-specific expression. Genes Development, October 2013.

94. A. Schep and S. University. motifmatchr: Fast Motif Matching in R, 2022.

95. Y. Hao, S. Hao, E. Andersen-Nissen, W. M. M. III, S. Zheng, A. Butler, M. J. Lee, A. J. Wilk, C. Darby, M. Zagar, P. Hoffman, M. Stoeckius, E. Papalexi, E. P. Mimitou, J. Jain, A. Srivastava, T. Stuart, L. B. Fleming, B. Yeung, A. J. Rogers, J. M. McElrath, C. A. Blish, R. Gottardo, P. Smibert, and R. Satija. Integrated analysis of multimodal single-cell data. Cell, 2021.

96. U. Raudvere, L. Kolberg, I. Kuzmin, T. Arak, P. Adler, H. Peterson, and J. Vilo. g:Profiler: a web server for functional enrichment analysis and conversions of gene lists (2019 update). Nucleic Acids Research, 47(W1):W191–W198, July 2019.

